# Effects of Reduced Watering Regimes on Hops Growth and Rhizospheric Microbial Communities

**DOI:** 10.64898/2026.11.03.764026

**Authors:** Alexandra McElwee-Adame, Caden Mulligan, Kailyn Kiburi, Zoe Glynn, Mani Golshani, Ahmed Alkhafaji, George Vourlitis, Arun Sethuraman

## Abstract

Marked by shifts in historic climatic patterns, the effects of climate change have begun to disrupt biological life worldwide. In agricultural systems, the abiotic effects of climate change such as increased temperatures and reduced rainfall extend beyond crop yield and can be felt at the ecological level through microbial community shifts associated at the interface of the root/soil matrix, known as the rhizosphere. Shifts in rhizospheric community composition have been known to greatly impact crop health, making common garden studies examining the totality of climate related impacts on crops of the utmost importance. Using a combination of both phenotypic and metabarcoding data collected from the rhizosphere, we present a multi-month drought tolerance experiment using the cash crop *Humulus lupulus* (hops) and five of its cultivars; Cascade, Centennial, Chinook, Comet and CTZ. Statistical analyses of phenotypic data revealed that hops displayed a differential impact on reduced watering strategies while rhizospheric data revealed more consistent patterns of association between microbes and cultivar type. We hope results from our study can be used to further expand our knowledge on the impact of future climate change and necessary reduced watering strategies on agricultural systems, providing insight into potential mechanisms of response for drought tolerance in crop systems.

**Highlight:** Impacts of reduced watering regimes varied significantly between cultivar types, revealing patterns in drought tolerance amongst cultivars with shared recent introgression events with USDA breeding wild male Utah 524 and 526.

## Introduction

Climate change has become one of the primary factors affecting agricultural production worldwide. Prior to the turn of the 21st century, research suggested that global temperatures would increase at a rate of nearly one third of a Fahrenheit degree each decade (Vose et al., 2021). However, studies in recent years have found that this increase in global temperatures is more likely to reach 1.5 °C or beyond by 2050 (Eftekhari, 2022), resulting in drastic alterations to global climate patterns such as changes in the frequency and intensity of extreme weather events such as floods and heatwaves, irregular precipitation patterns, and reduced snowpack leading to decreased water availability during growing seasons (Arora, 2019; Malhi et al., 2021). At the organismal level, these abiotic stressors can cause damage to crop physiology by either physically damaging the crop itself, therefore directly reducing product yield (Bibi & Rahman, 2023; Yuan et al., 2024) or by altering the rhizospheric microbiome and biogeochemistry of the surrounding soil (Adebayo et al., 2026).

Many crop species are known to harbor beneficial microbes known as plant-growth-promoting microorganisms (PGPM) that assist in nutrient absorption key for proper growth and development as well as water-use efficiency. Disturbances to the plant-surrounding microbiome can result in shifts in the composition and abundance of the microbial community (Laishram et al., 2024; Muhammad et al., 2025). At the cellular level, microbes classified as PGPM’s have been found to accumulate a surplus of vital compounds such as amino acids and sugars, which are necessary to withstand degradation caused by unfavorable or infrequent watering conditions (Jha & Saraf, 2015). Bacterial genera such as *Bacillus* have been of high interest to crop breeders due to their associations being marked with an increase in resistance to drought in several major crops, such as maize (Vardharajula et al., 2011), peppers (Lim & Kim, 2013), soybean (Sheteiwy et al., 2021), wheat (Rashid et al., 2022; Sood et al., 2020), chickpea (Kumar et al., 2016) and cannabis (Wiszpolski & Stolarski, 2026). Therefore, in order for plant breeders to properly prepare and develop new and more resilient cultivated crop varieties in the future, a holistic approach is needed to understand the effects of abiotic stress on the totality of the plant itself and reveal underlying mechanisms that may be involved in abiotic stress tolerance.

For crops produced in the mid- and high-latitude ranges, ideal growing ranges in future years will need to expand northward (King et al., 2018); however, for some farmers, especially those in northern regions like the Pacific Northwest of the United States, this may not be an option. Changes in precipitation regimes in particular are expected to strongly impact farmers at regional levels, resulting in inconsistent water availability (Feng et al., 2019; Konapala et al., 2020). As a result, farmers will need to turn to selective plant breeding strategies to address consequences associated with climate change in these regions.

Hops (*Humulus lupulus*) is a dioecious perennial cash crop that is natively distributed throughout the Eurasian and North American continents; however, through breeding program efforts, hops can now be produced globally (Murakami et al., 2006). Hops display an ideal growing range between 45° and 50° north of the equator but have become locally adapted to various regions outside of this range via selective breeding efforts (Castro et al., 2021; Ruggeri et al., 2024). Commercially, over 250 cultivars (cultivated varieties) have been developed worldwide, each displaying differences in bitter acid and essential oil compositions, key benchmarks in the brewing industry (McElwee-Adame et al., 2025), as well as differences in agronomic traits such as increased growth, yield, seasonality, and disease/pest tolerance (McAdam et al., 2013).

Majority of the global hops production is dominated by two countries: Germany and the United States (Agraso-Otero et al., 2026). Unfortunately, in recent years, US production has seen a decline in acres of hops harvested and value of production, with the 2023 growing year recording a devastating 21% decline in overall production value compared to the previous growing year (USDA-SASS National Hops Report, 2024). Coincidentally, in recent years, the Pacific Northwest (PNW) has seen extreme changes in climatic regimes, with increased summer temperatures and drought events becoming more common across the region.

Previous studies utilizing simulated data have found that higher global temperatures are also expected to decrease overall hops production and alter bitter acid production and essential oil compositions (Mozny et al., 2009, 2023). Of the total hops produced worldwide, an overwhelming majority (∼98%) is used by the brewing industry (Korpelainen & Pietiläinen, 2021), valued at 113 billion dollars (US) in the United States beer market alone (Brewers Association, 2025), meaning that alterations to future hops products will directly alter future beer production. Additionally, previous studies aiming to generate real-time data using common garden experiments have already begun to assess the impact that reduced watering regimes and increased temperatures may have on hops growth and production (Eriksen et al., 2020; Nakawuka et al., 2017). However, both these studies evaluated the impact of abiotic stress on hops solely from a physiological standpoint, completely negating belowground associations.

Here we present a multi-month common garden experiment addressing the impact of reduced watering regime strategies on hop growth and rhizospheric microbial associations. By employing a combination of genomic and phenotypic data, we address three main questions: 1) Which of the selected cultivars is least impacted by drought, therefore making it an ideal candidate for future breeding program efforts? 2) Are there any patterns between hops’ resilience under reduced watering strategies and pedigree? 3) Could belowground associations such as rhizospheric microbial and fungal compositions confer drought tolerance? We hypothesize (1) that cultivars that share a more recent hybridization event with wild American hops will display a higher tolerance for drought via adaptive introgression, (2) that this pattern of tolerance will be shared amongst individuals of similar pedigree history, and (3) elevated community composition of known beneficial microbes coevolved in response to local adaptation to North American soils.

## Methods

### Cultivar Selection and Pre-Drought Regime Care

Five cultivated varieties were selected based on the 2023 National Hops Report based on two criteria: top-producing cultivars during the 2023 growing season and access to rhizomes. The following cultivars were then selected: Cascade (USDA Accession #: 56013), Centennial USDA Accession #: 21507), Chinook (USDA Accession #: 21226), Comet (USDA Accession #: 62013), and Columbus/ Tomahawk/ Zeus (hereon, CTZ). In April 2024, 150 rhizomes per cultivar (total n=750) were purchased from The Thyme Garden nursery (Alesca, Oregon) and shipped to San Diego State University (SDSU), stored at 4C until transport to maintain dormancy. Rhizomes were then transported from SDSU to the California State University San Marcos (CSUSM) Biology Department Greenhouse (33.131294, -117.156819) and planted roughly two inches below the soil line into one gallon nursery pots with potting soil composed of aged forest product, sphagnum peat moss, perlite and 0.3-0.3-0.5 NPK fertilizer derived from earthworm castings, bat guano, oyster shell, and dolomite lime (FoxFarm Soil & Fertilizer Co 2026). No additional amendments or fertilizers were added to the experimental pots for the remainder of the experiment.

From April 2024 to June 2025, juvenile plants were reared at the CSUSM greenhouse. To minimize the risk of root rot amongst our juvenile individuals, plants were watered twice a week via manual irrigation from a hose, achieving roughly 1 inch of water within each 1-gallon pot per watering session from April 2024 to October 2024, and then once a week from October 2024 to the end of June 2025. At the end of June 2025, 90 individuals per cultivar were randomly selected (n=450) and assigned one of three treatment regimes: a control treatment (100% recommended watering), moderate drought (75% recommended watering), and severe drought (50% recommended watering). Plants were then transferred to SDSU’s Alvarado Greenhouse (32.7771272, -117.0629152) for the twelve-week drought experimental period (7/1/25-9/23/25). Due to low germination success with our Chinook and Comet rhizomes, 90 Chinook and 60 Comet juvenile plants were purchased from Great Lake Hops (Ottawa County, MI) and used for the experimental period.

### Drought Regime Delivery and Experimental Design

To standardize delivery of water regimes, we constructed three drip irrigation systems (one per treatment) and utilized a timer system for regime delivery. Individuals of the same treatment type (n=30 per cultivar type; n= 150) were randomly placed along a greenhouse bench and were attached to one of three irrigation systems: control, moderate, or severe. For our experiment, we opted to establish our control conditions as receiving 1.5 inches of water per week. We measured the volume necessary to create 1.5 inches of water within our nursery pots, which equated to roughly 850 mL/week of water for our control treatment. From there, we calculated that our moderate regime individuals would receive 600mL/week (75% recommendation) and the severe regime would receive 425mL/week (50% recommendation).

To avoid oversaturating the root system of our hop plants, we opted to split weekly waterings into two disbursements per week, resulting in individuals receiving either 425 mL (control), 300 mL (moderate), and 215 mL (severe) per disbursement. To ensure that each individual within each treatment type received the same amount of water, we timed each treatment system to estimate the amount of time it took to achieve each of our targeted volumes (425 mL, 300 mL, and 215 mL) for each disbursement, resulting in about 7 minutes of watering for our control treatment, 5 minutes for moderate, and 3 minutes for severe treatment. Randomized spot checks were performed throughout the duration of the experiment to monitor for consistency of water delivery along irrigation lines.

Daily environmental conditions such as hourly average temperature and relative humidity were recorded and monitored via HOBO data loggers (Onset Computer Corporation, Bourne, MA). During the 12-week drought period, plants received an average of 12-14 hours of daylight per day. Additionally, plants during the experimental period were maintained and cared for through weekly maintenance such as trimming dead shoots or vines to promote growth and removal of agricultural pests such as spider mites via predation by the biocontrol agent *Hippodamia convergens* (via NaturesGoodGuys, <u>Amazon.com</u>) and were released twice per growing season.

### Phenotypic Data Collection

To measure each cultivar’s growth rate response to varying levels of drought, weekly vertical height was measured for each individual, taken from the base of the plant along the soil line to the tip of the apical shoot. In the event that the apical shoot was not available (i.e., died off due to heat stress), we would then use the next available lateral shoot to serve as a proxy apical shoot for measurements. Vertical growth measurements were recorded weekly over the twelve-week growing season. At the end of the twelve-week growing season, weekly growth rates were calculated from our data using the following equation: h_t+1_ - h_t_ where t is the week of interest and h is the vertical height for that particular week. Additionally, at the end of the experimental period, aboveground biomass was collected for each individual by trimming plants down to the base of the soil line, bagged, and weight recorded. Leaf samples for each individual were then bagged and stored in -80C freezer for downstream analyses.

### Difference in Biomass Production and Average Growth Rate (AGR)

To test hypotheses of the impacts of cultivars or watering regimes on weekly growth rates or biomass production, an ANOVA was performed using the base R function aov() (R Core Team). For these ANOVA analyses, treatment type, cultivar type and week were treated as the independent variables while weekly growth rates and biomass were treated as the dependent or response variables. First, we assessed each watering regime dataset (control, moderate, and severe) for normality and homoscedasticity using the base R functions shapiro.test() and bartlett.test(). If both assumptions were met, a Tukey Post Hoc test was performed on statistically significant datasets to identify significant pairwise differences using the base R function TukeyHSD(). If a dataset failed to meet normality, the R package best.normalized (Peterson & Cavanaugh, 2020) was used to determine and perform the optimal data transformation for either biomass or weekly growth rate measurements. Tests for normality and homoscedasticity were then repeated, and if assumptions were met, a Tukey Post Hoc test was performed as described above. Significance was assessed using the threshold adjusted p-value < 0.05, assuming a null hypothesis of no significant difference in biomass between cultivar or treatment types. If datasets failed ANOVA assumptions, we opted to use a Kruskal-Wallis test using the kruskal.test() base R function followed by a Dunn test for post hoc testing using the rstatix package (Kassambara, 2026).

To test hypotheses of the impacts on weekly growth rates across cultivars and treatment types, we performed a linear mixed-effects model (LMM) fitted by a restricted maximum likelihood (REML) method using the R packages lme4 (Bates et al., 2015) and lmerTest (Kuznetsova et al., 2017). Week, treatment, cultivar, and treatment∼cultivar were treated as fixed effects, and growth as the response variable. Next, we performed a Type III ANOVA with Satterthwaite’s method with outputs from our LMM using the base R function aov(). Lastly, least-squares means calculated from our LMM were used in downstream pairwise comparisons using the emmeans R package (Lenth and Piaskowski 2026), adjusting for multiple comparisons using a Tukey adjustment. Additionally, both biomass and growth rate data were visualized using ggplot2 (Wickham 2016).

### Measurements of Physiological Responses to Drought

To test hypotheses of leaf physiological responses to drought, we measured leaf surface area and circularity at the end of the 12-week growing season across all three treatment types. Two leaf samples per individual (n=28) were collected from randomly selected plants and photographed. Photographed images were scaled by a standardized reference and converted to JPEGs. JPEG files were then uploaded into the software ImageJ (Schneider et al., 2012), and each leaf was manually filled with a solid red overlay (RGB, 255, 0, 0) to create a region of interest (ROI) image. Thereon, ImageJ was utilized to calculate a proxy for leaf surface area using the number of pixels found within the ROI and leaf perimeter, using the border established by the ROI. Lastly, circularity was calculated using the following equation by Hightower et al., 2024: (4π ×Area)/Perimeter^2. Leaf surface areas and circularity were averaged across the two leaves per individual per cultivar per treatment. Differences between cultivar or treatment types’ leaf surface area and circularity values were then statistically tested using the base aov() function in R, verified to meet standard ANOVA assumptions, and assessed for significance using a Tukey Post Hot test with a p-value threshold of 0.05.

### Rhizospheric Collections and Sequencing

To explore the real-time effects of drought on hops throughout their lifecycle during the growing season, we collected soil material at two time points: Week 2 and Week 12. We opted to collect soil material at Week 2 to provide plants and their respective rhizospheric biomes the time to acclimate to their new watering regime, and to explore both the immediate and long-term effects of reduced watering regimes on the plants. Ten individuals per cultivar for each treatment type were randomly selected for leaf and soil collection at both time points (n= 300). Soil material adjacent to roots was collected two inches below the surface, scraping any present roots available. Soil material was then stored on ice and placed in the -80C freezer for storage until use.

Two individuals per cultivar per treatment were then randomly selected for downstream sequencing (n=30). 0.5g of soil material collected from both Week 2 and Week 12 was loaded into individual microcentrifuge tubes for DNA extraction, library prep, sequencing, and taxon identification/quantification all performed by Zymo Research (Irvine, CA). For this study, we opted to utilize both 16S and ITS sequencing to explore the entirety of the rhizospheric diversity in microbial and fungal biomes. Briefly, gDNA was extracted and purified from collected soil samples using the ZymoBIOMICS^®^-96 MagBead DNA Kit for 16S sequencing. Next, targeted genes were amplified from the V3-V4 region of the 16S rRNA gene for 16S sequencing and the ITS2 region for ITS sequencing using the *Quick*-16S^™^ NGS Library Prep Kit (Zymo Research, Irvine, CA) and amplified via RT-PCR. Amplified fragments were then sequenced on Illumina^®^ Nextseq^™^ with a P1 reagent kit for 600 cycles.

Raw reads generated from sequencing were then analyzed using the DADA2 pipeline (Callahan et al., 2016) to remove potential sequencing errors and chimeric sequences and to identify unique variants. Variants were then used to infer operational taxonomic units (OTUs) and OTU compositions per individual using Uclust from QIIME v.1.9.1 (Caporaso et al., 2010). Identified OTUs and percent abundance were then downloaded into a CSV file and used for downstream statistical analysis.

### Patterns in Rhizospheric Diversity

To test hypotheses of the effects of treatment, duration of drought (start vs end), or cultivar types on microbial abundance, we performed a linear mixed model test (LMM). Due to the amount of microbial diversity discovered from 16S and ITS sequencing, we split our OTU dataset into three groups: fungal (n=85), archaeal (n=6), and bacterial (n=174). We opted to perform these LMMs separately rather than together to reduce model complexity and to reduce computational complexity. For each microbial/fungal OTU data subset, an LMM was performed using treatment, timepoint, and cultivar as the fixed effects and percent abundance within the rhizosphere as the response variable and individual as the random effect. Bacterial, archaeal, and fungal species identified as highly significant (p-value < 0.001) were then explored in the literature to identify previously known drought tolerance associations.

### Analysis of Rhizospheric Nutrient Bioavailability

Soil was analyzed for extractable phosphate (PO4), nitrogen (nitrate and ammonium), and dissolved organic carbon (C) to assess how P, N, and C were altered by hops varieties and drought treatments. Extractable P was analyzed using the ascorbic acid method (Kuo 1996). Briefly, approximately 5 g of fresh soil was added to 50 mL of a 0.05*M* HCl + 0.0125*M* H2SO4 extraction solution and shaken on a table shaker (E-Class, Barnstead/Lab-Line, Dubuque, IA) for 10 minutes. Extracts were filtered using qualitative filter paper (P8, Fisher Scientific, Inc., Pittsburg, PN), and the extract was measured at 880 nm after adding a molybdate color reagent and an ascorbic acid reducing solution. Extractable N and dissolved organic C (DOC) were measured from 0.5*M* K2SO4 extracts where approximately 10 g of fresh soil was extracted in 40 mL 0.5*M* K2SO4 after shaking on a table shaker for 60 minutes. For extractable N, filtered extracts are analyzed for NH_4_ (Mulvaney 1996) and NO_3_ (Miranda et al. 2001) concentration by absorbance at 667 nm (NH_4_) and 540 nm (NO_3_) using a spectrophotometer (Spectroquant Pharo 300, Millipore Sigma, Burlington, MA, USA). For DOC, extracts were digested using the Walkley-Black method (Sims and Haby 1971) and measured colorimetrically at an absorbance of 600 nm with a spectrophotometer (Spectroquant Pharo 300, Millipore Sigma, Burlington, MA, USA). Data are presented as concentrations per unit soil dry weight after correcting for the moisture content of fresh soil. Dissolved inorganic N is defined as the sum of extractable NH4 + NO3.

## Results

### Impact on Biomass Production

To assess whether reduced watering strategies impacted hops aboveground biomass production over the 12-week experimental period, we performed either a two-way ANOVA test (Cascade, Centennial, Chinook and Comet) or a non-parametric Kruskal-Wallis test (CTZ) on collected biomass values (g). Cascade (df= 2, F= 0.064, p= 0.939), Centennial (df= 2, F= 0.918, p= 0.403), and Comet (df= 2, F=2.899, p= 0.067) showed no significant difference in biomass production between control, moderate, and severe treatment types (Figure 1; Table 2). Chinook’s (df=2, F= 21.020, p= 7.98E-07) average biomass was the highest under moderate treatment type and lowest under the control treatment (Table 1; Figure 1). Biomass production significantly differed between all treatment types, with our Tukey post-hoc test revealing moderate versus control treatments displaying the greatest impact (Table 3).

**Fig 1.**
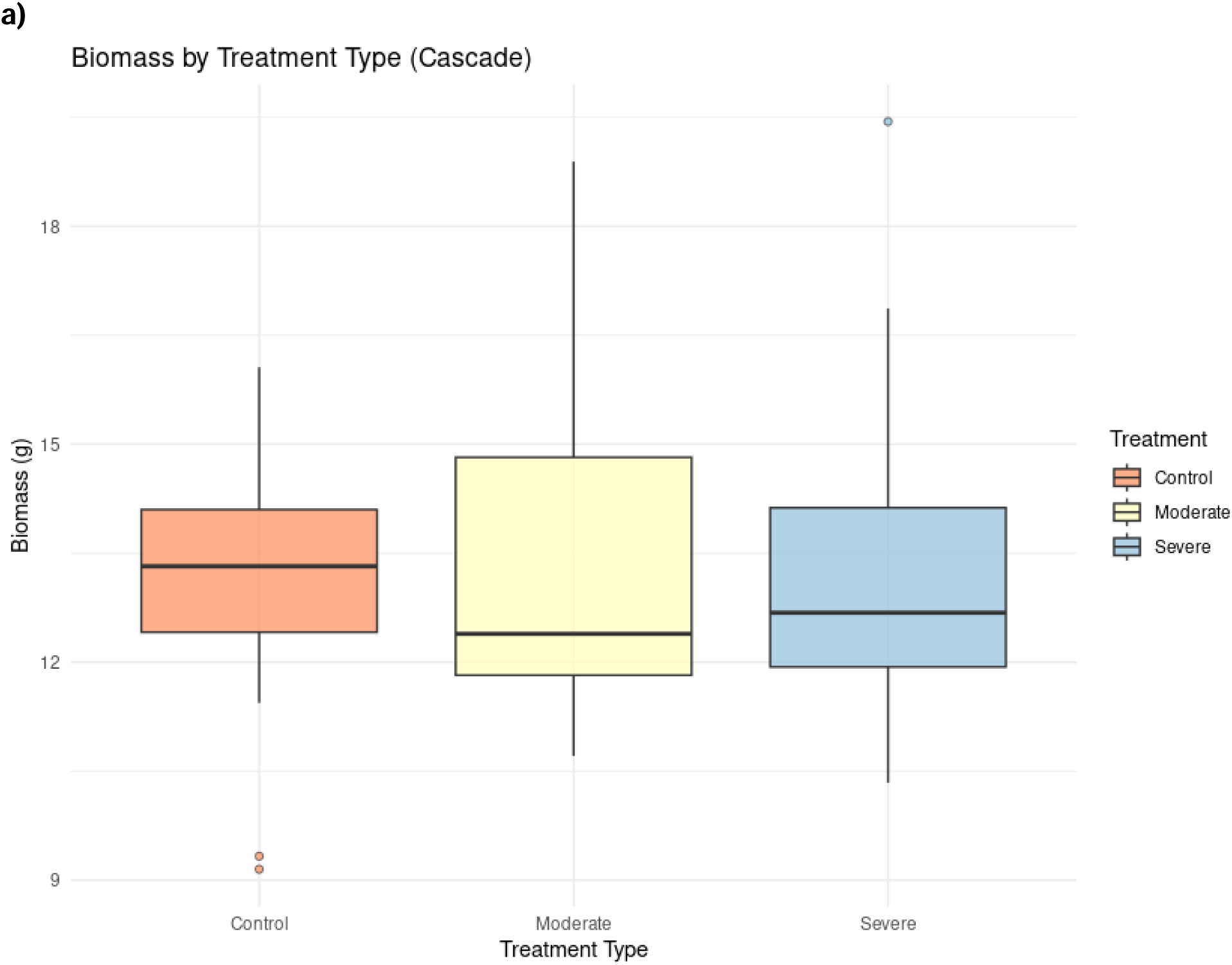
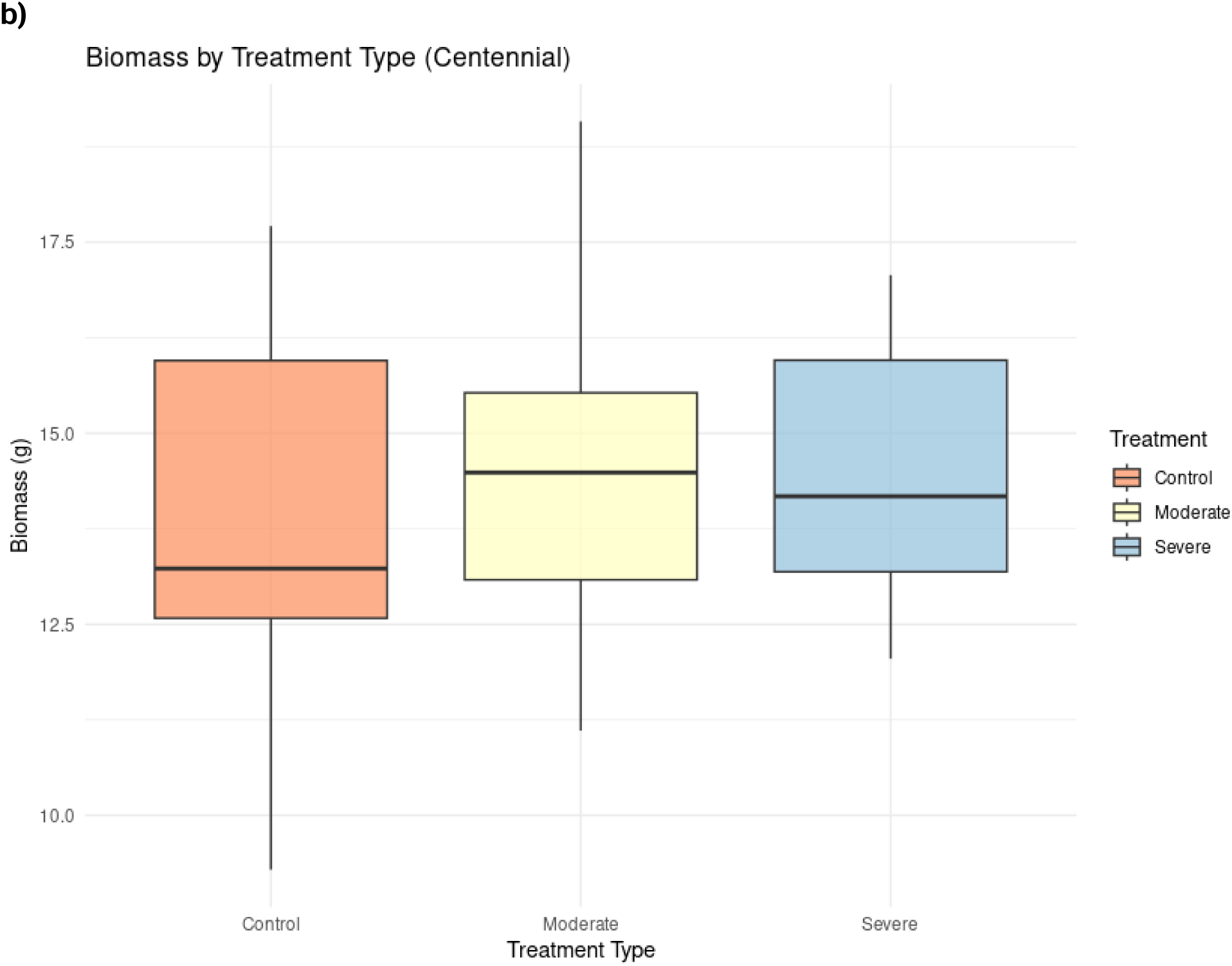
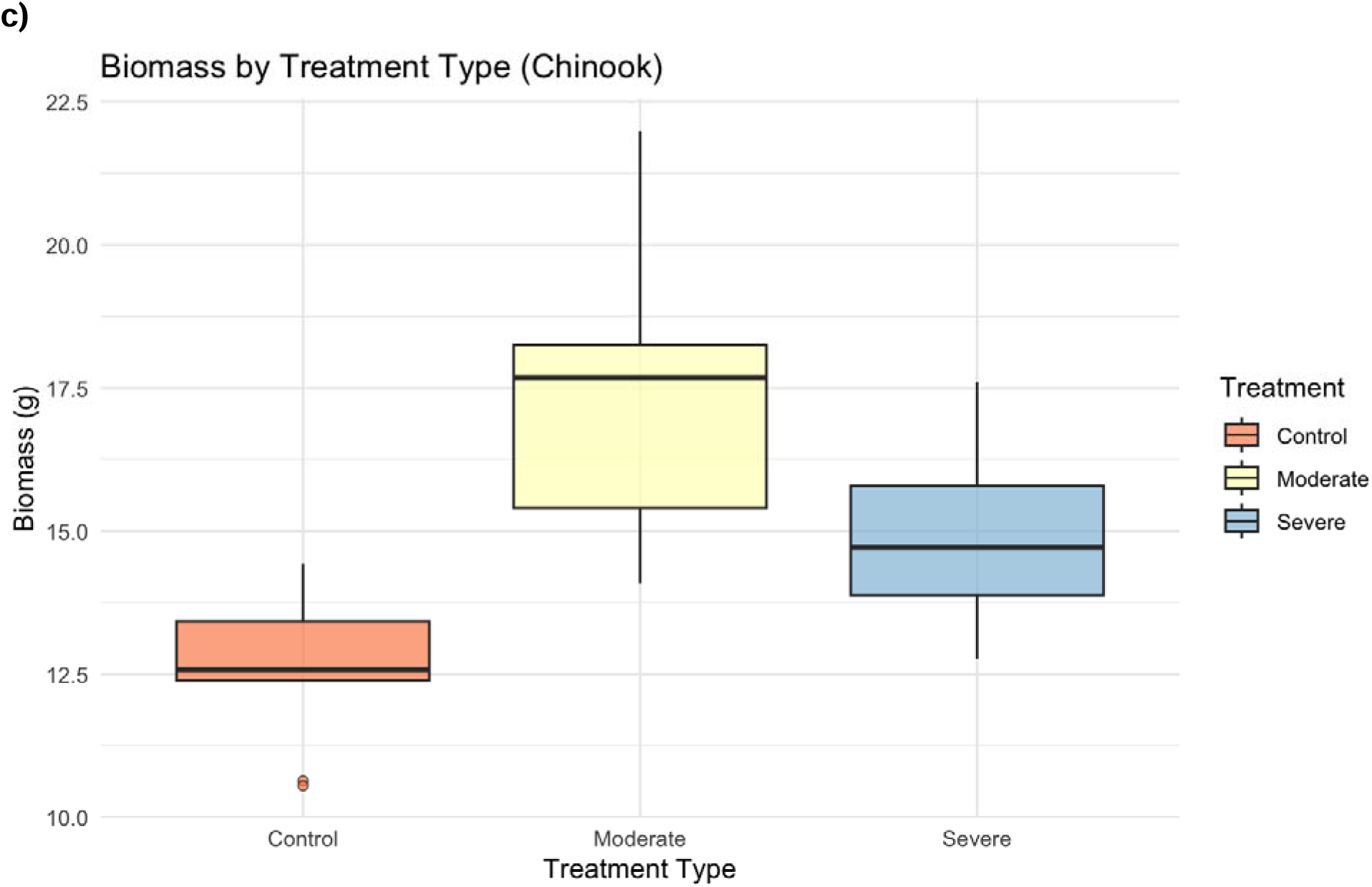
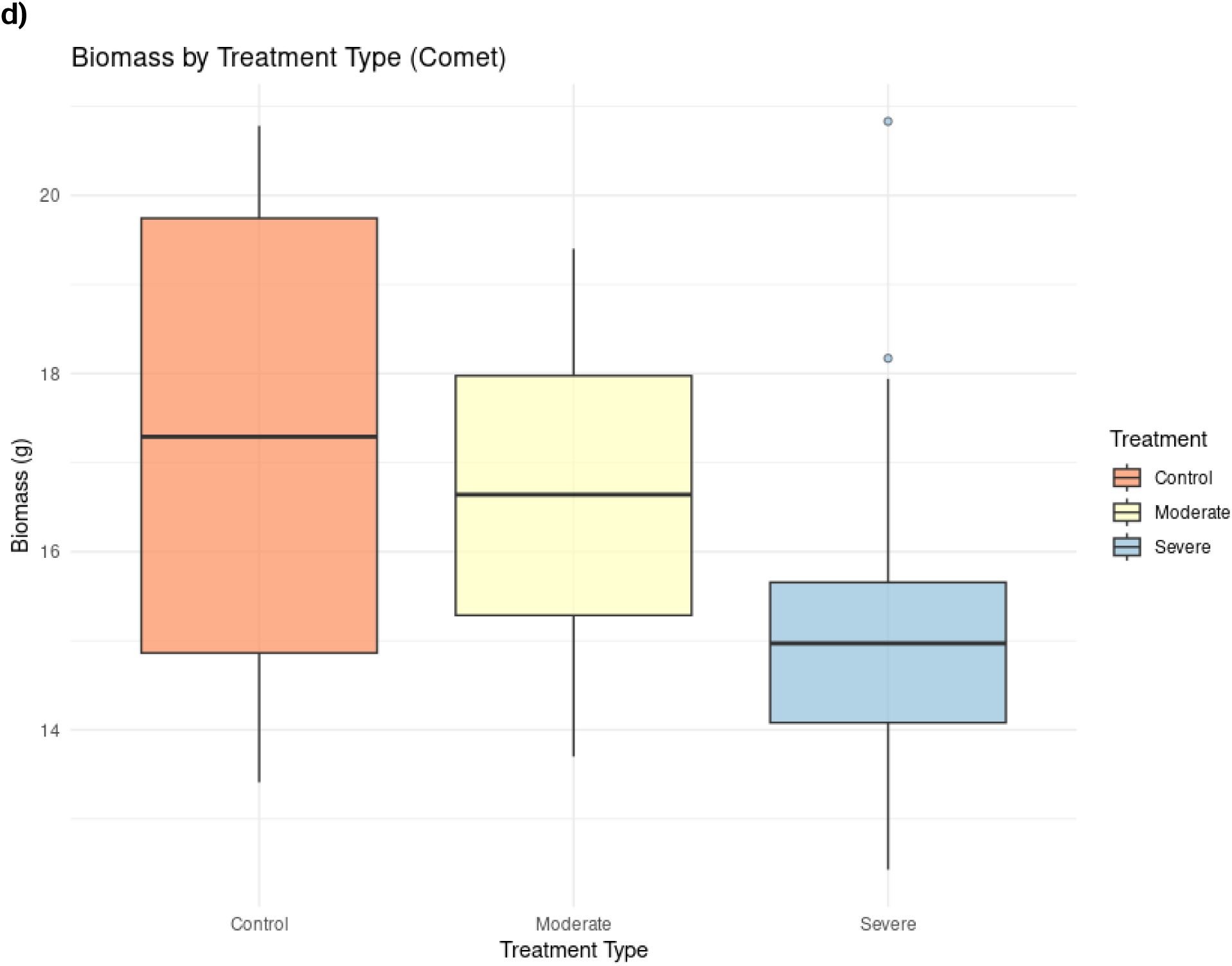
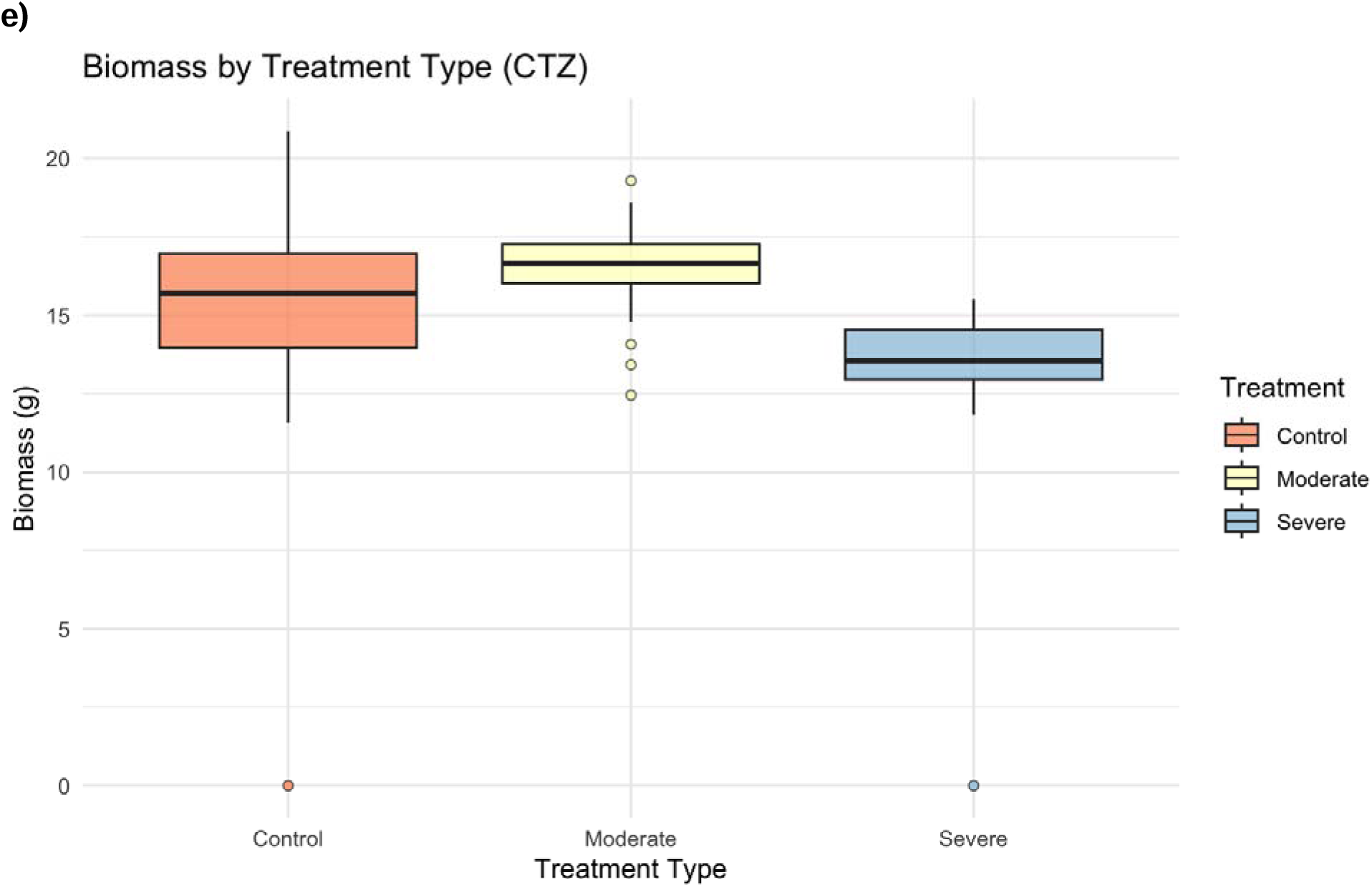
Boxplots of measured aboveground biomass (g) for Cascade (a), Centennial (b), Chinook (c), Comet (d) and CTZ (e) across the three treatment types control (red), moderate (yellow) and severe (blue).

**Table 1.**
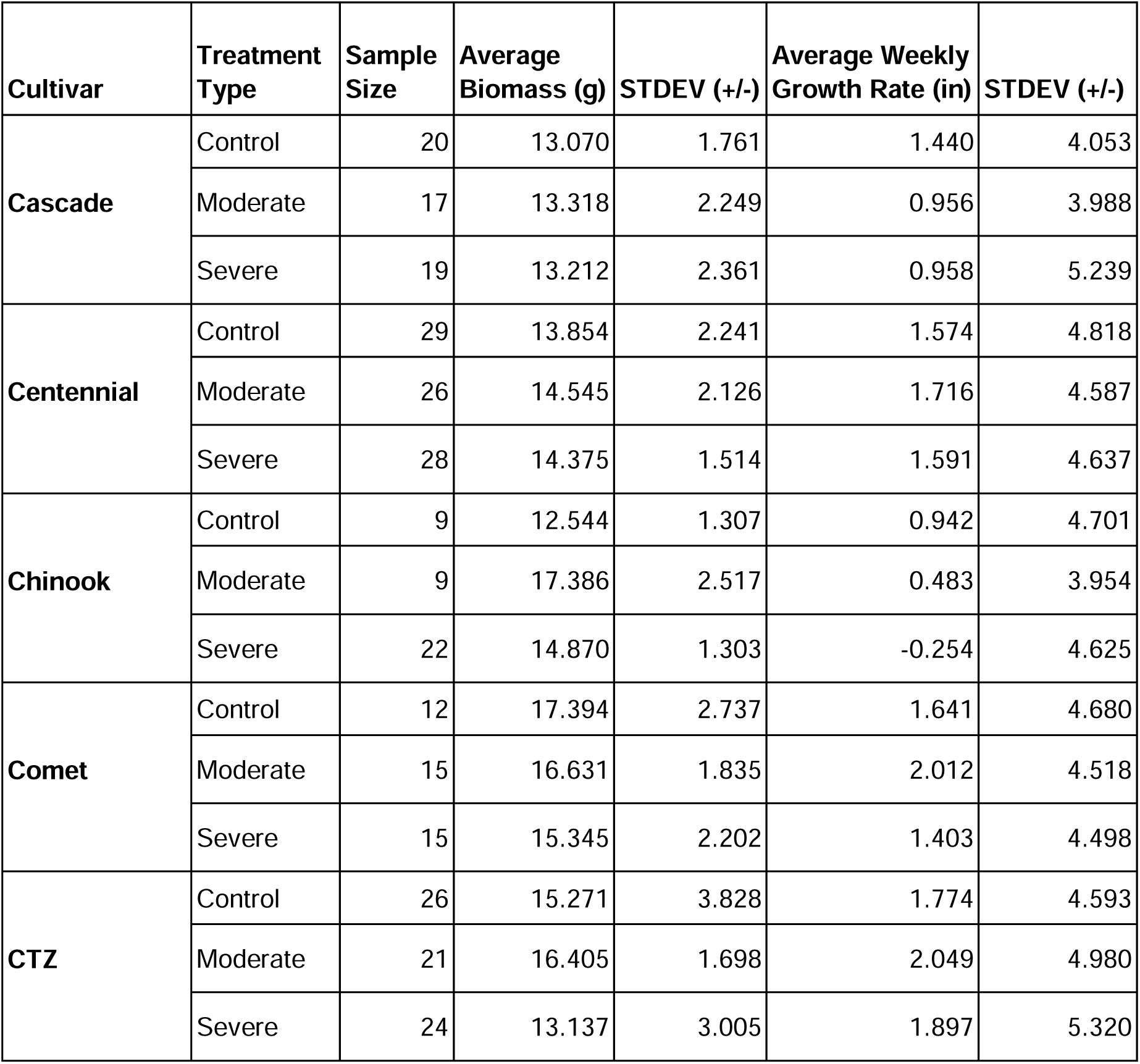
Table of sample size, average biomass, and weekly growth rates for each cultivar type across their various treatments.

**Table 2.**
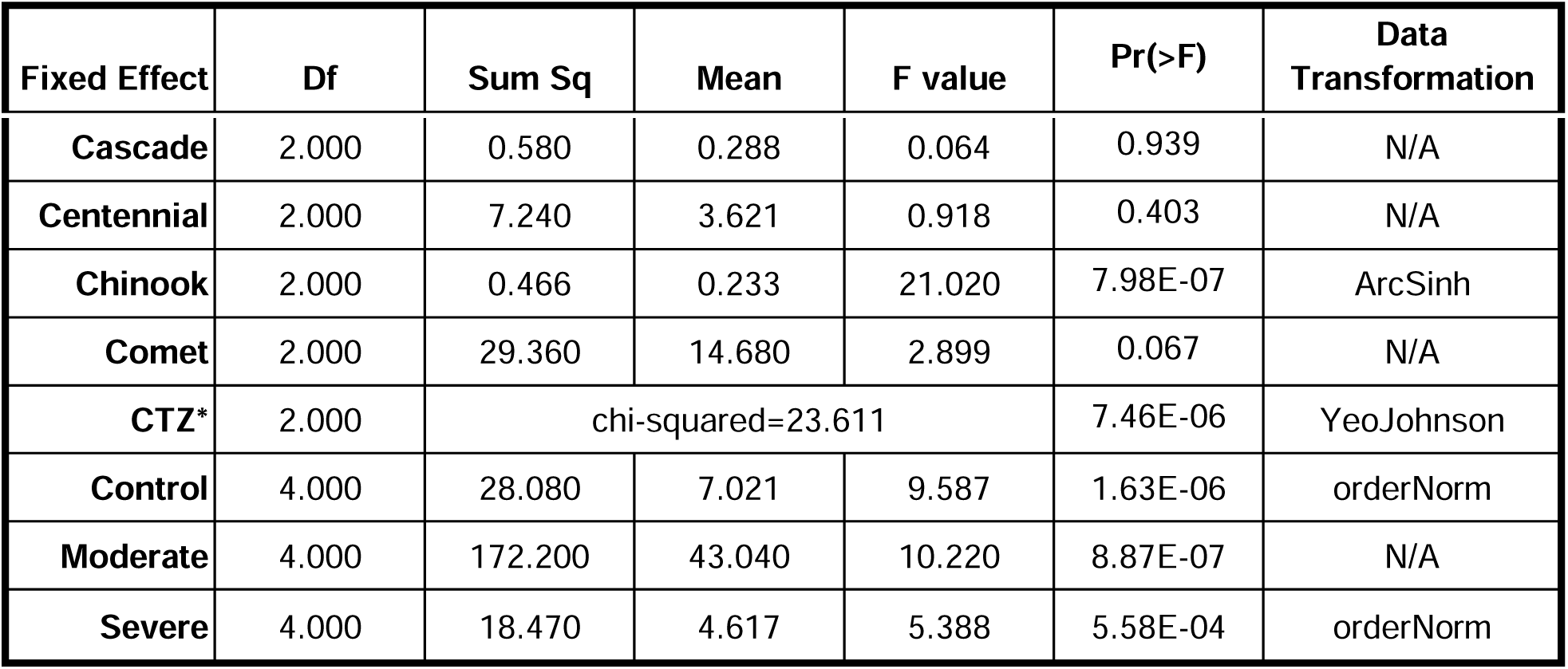
Summary statistics and p-values generated from ANOVA test for aboveground biomass values (g). Fixed effects tested: cultivar type (Cascade, Centennial, Chinook, Comet, CTZ) and treatment type (Control, Moderate, and Severe) with aboveground biomass as the response variable. * indicates the use of a Kruskal-Wallis test in lieu of an ANOVA test when necessary.

| **Fixed Effect** | **Df** | **Sum Sq** | **Mean** | **F value** | **Pr(>F)** | **Data Transformation** |
|---|---|---|---|---|---|---|
| **Cascade** | 2.000 | 0.580 | 0.288 | 0.064 | 0.939 | N/A |
| **Centennial** | 2.000 | 7.240 | 3.621 | 0.918 | 0.403 | N/A |
| **Chinook** | 2.000 | 0.466 | 0.233 | 21.020 | 7.98E-07 | ArcSinh |
| **Comet** | 2.000 | 29.360 | 14.680 | 2.899 | 0.067 | N/A |
| **CTZ<sup>\*</sup>** | 2.000 | chi-squared=23.611 |  |  | 7.46E-06 | YeoJohnson |
| **Control** | 4.000 | 28.080 | 7.021 | 9.587 | 1.63E-06 | orderNorm |
| **Moderate** | 4.000 | 172.200 | 43.040 | 10.220 | 8.87E-07 | N/A |
| **Severe** | 4.000 | 18.470 | 4.617 | 5.388 | 5.58E-04 | orderNorm |

**Table 3.** Generated p-values from post-hoc (Tukey or Dunn) testing between treatment types for each cultivar type using biomass as the response variable.

|  | **Moderate~Control** | **Severe~Control** | **Severe~Moderate** |
|---|---|---|---|
| **Cascade** | 0.934 | 0.976 | 0.988 |
| **Centennial** | 0.406 | 0.585 | 0.947 |
| **Chinook** | 4.00E-07 | 6.17E-04 | 2.52E-03 |
| **Comet** | 0.658 | 0.060 | 0.273 |
| **CTZ** | 0.164 | 9.95E-04 | 9.27E-06 |

A non-parametric Kruskal-Wallis test on CTZ’s biomass data (due to failed parametric ANOVA assumptions) revealed that biomass production significantly differed between treatment types (df=2, x^2^= 23.611, p= 7.46E-06) (Table 2). A post hoc Dunn test revealed significant differences between control versus severe (p= 9.95E-04) and moderate versus severe (p= 9.27E-06) models (Table 3), with no significant difference in biomass production between control and moderate (p= 0.164) regimes.

When treatment type was used as a fixed effect, aboveground biomass production varied significantly for all three treatment types; control (df= 4, F= 9.587, p= 1.63E-06), moderate drought (df= 4, F= 10.22, p=8.87E-07) and severe drought (df= 4, F= 5.388, p=5.58E-04). Tukey post-hoc testing revealed that under the control treatment, Comet and CTZ produced significantly larger (p < 0.05) aboveground biomass than Cascade (∼Comet, p=5.67E-05; ∼CTZ, p= 3.84E-03), Centennial (∼Comet, p= 6.84E-04; ∼CTZ, p= 0.048), and Chinook (∼Comet, p=1.32E-04; ∼CTZ, p= 6.17E-03) (Figure 2; Table 4). Under the moderate drought treatment, Chinook displayed insignificant differences in aboveground biomass when compared to Comet (p= 0.906) and CTZ (p= 0.751). Under the severe treatment type, aboveground biomass most noticeably differed between Comet and two other cultivars, with Comet displaying significant impacts on biomass production compared to Cascade (p= 4.19E-03) and CTZ (p= 0.021), but insignificant differences with Centennial (p= 0.537) and Chinook (p= 0.979) (Figure 2; Table 4).

**Fig 2.**
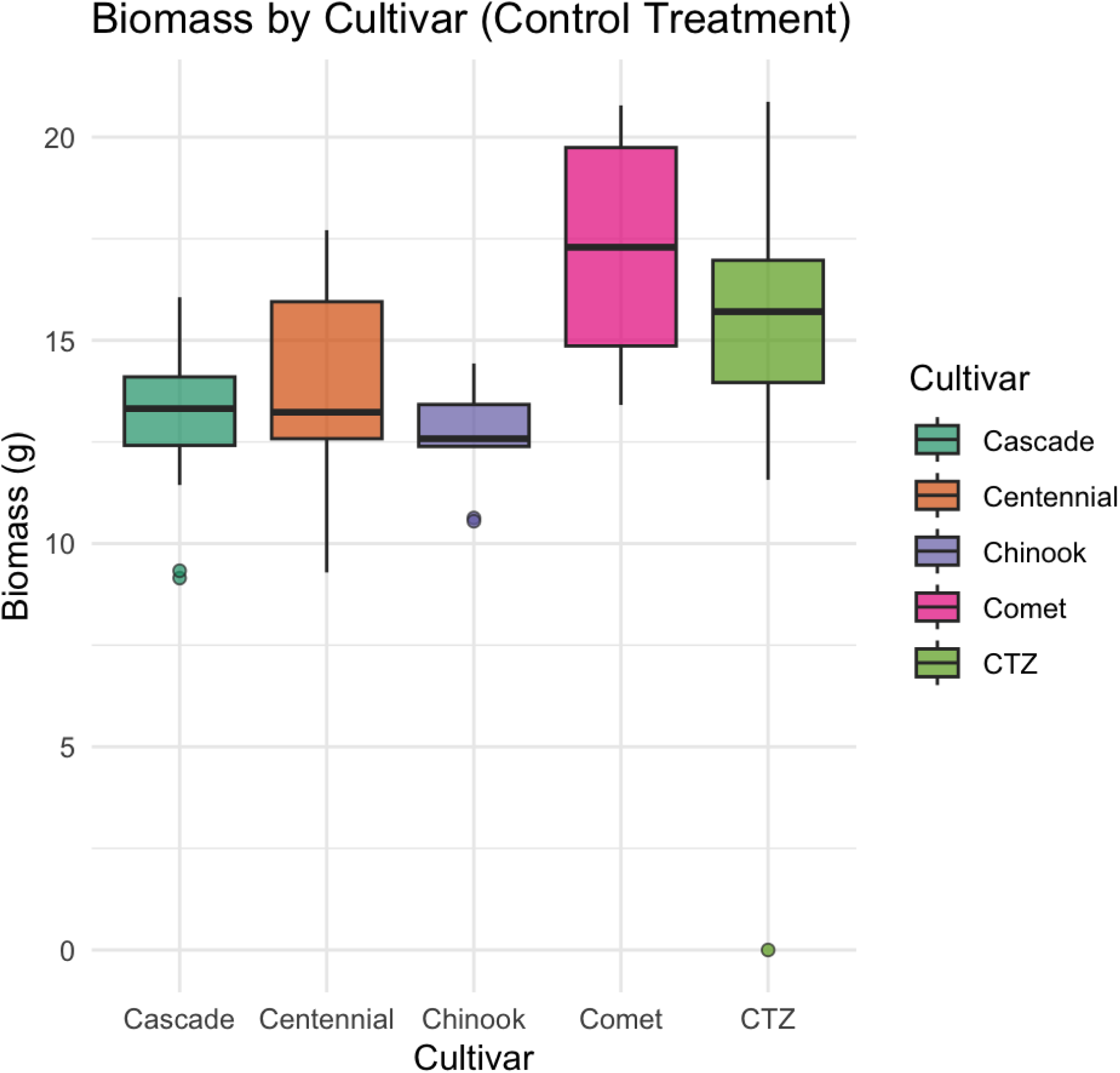
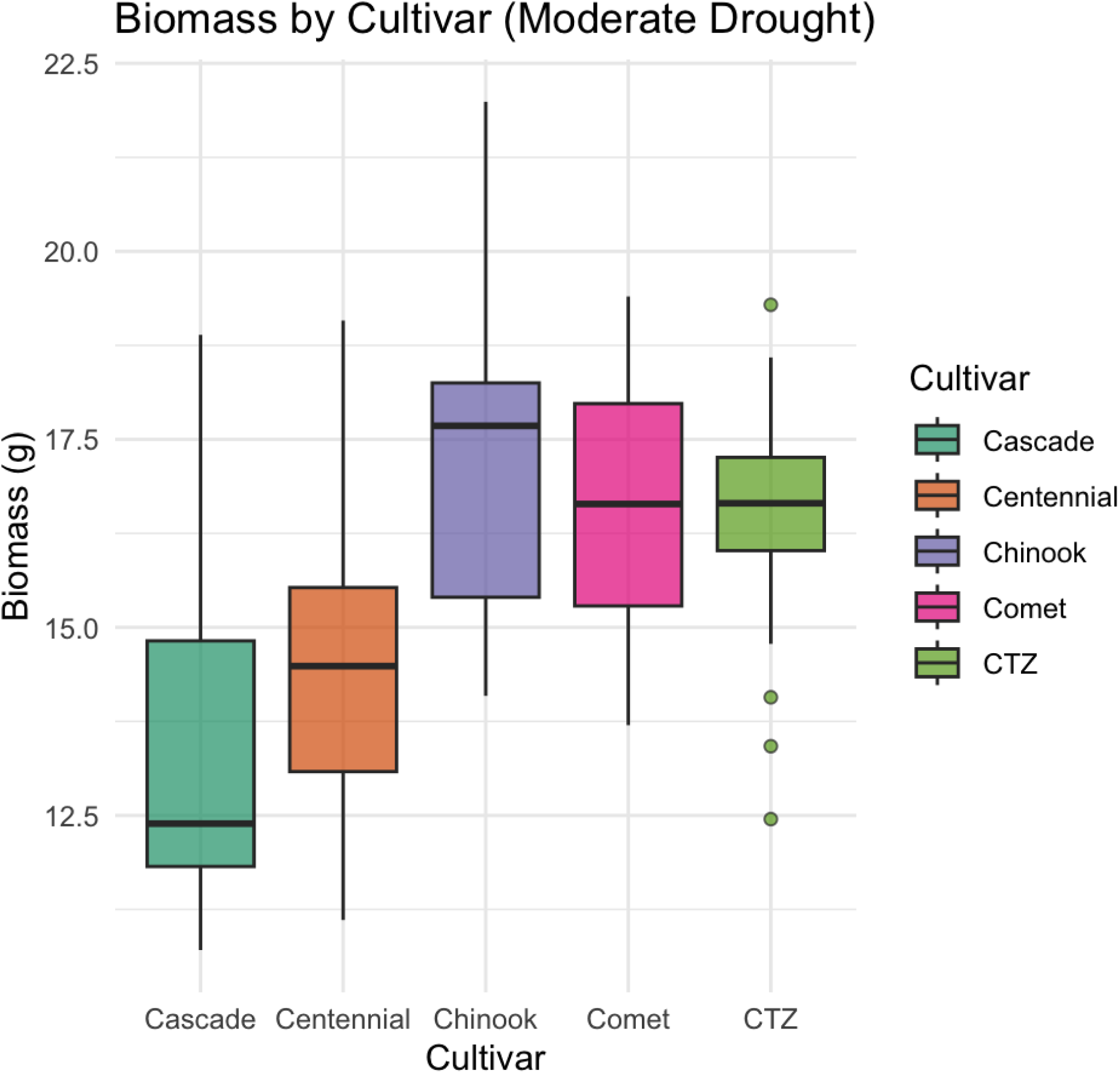
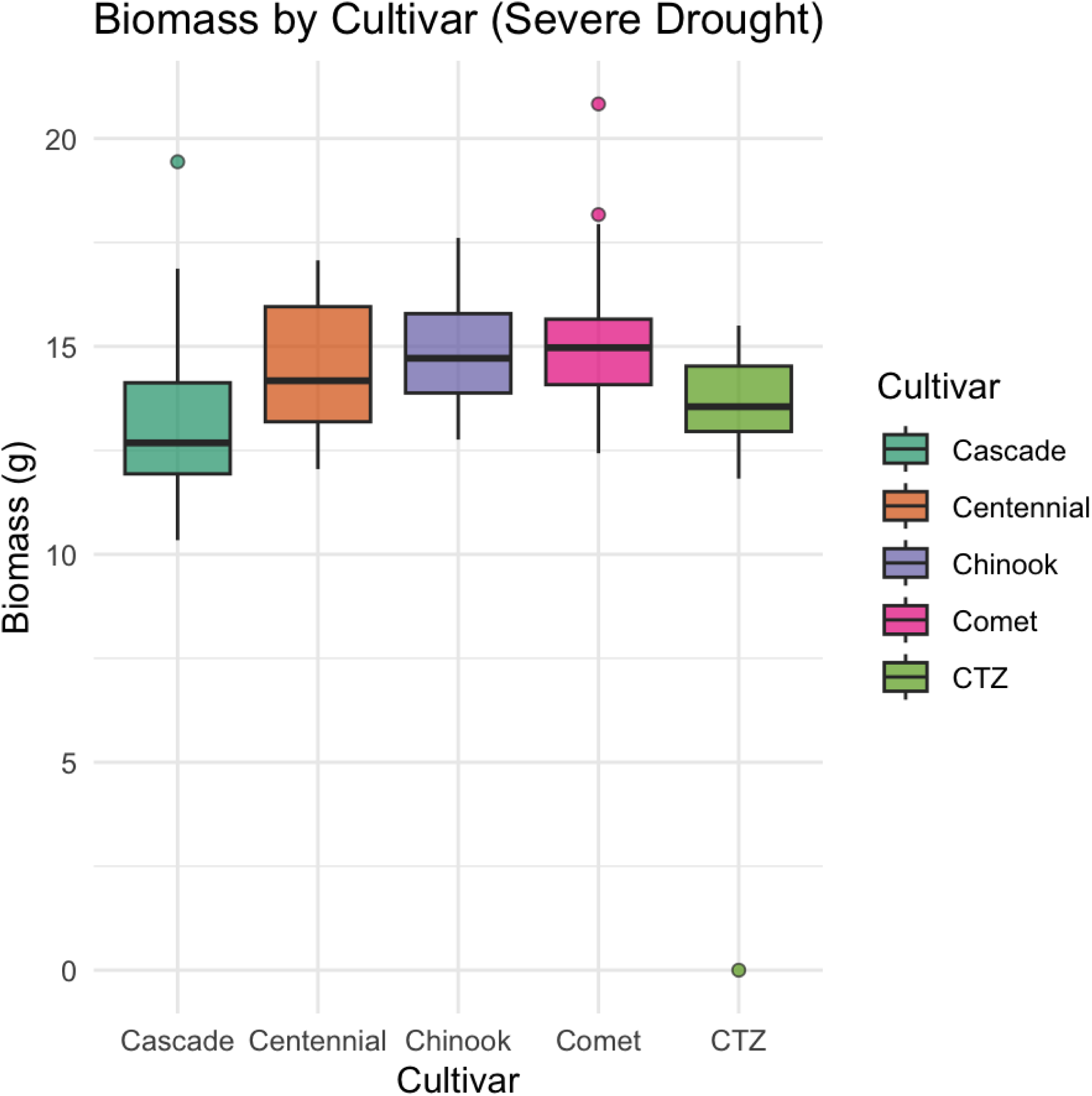
Boxplots of aboveground biomass production in grams, across cultivars within watering regimes - (a) Control, (b) moderate drought, and (c) severe drought Teal= Cascade, orange= Centennial, purple= Chinook, pink= Comet and green= CTZ.

**Table 4.** p-values generated from Tukey Post Hoc testing of ANOVA results using treatment type as the fixed effect and aboveground biomass as the response variable.

|  | Cascade | Centennial | Chinook | Comet |  |
|---|---|---|---|---|---|
| **Centennial** | 0.7850847 | - | - | - | **Control Treatment** |
| **Chinook** | 0.9619241 | 0.5170894 | - | - |  |
| **Comet** | 5.67E-05 | 6.84E-04 | 1.32E-04 | - |  |
| **CTZ** | 3.84E-03 | 0.0480085 | 6.17E-03 | 0.3134526 |  |
| **Centennial** | 0.3165009 | - | - | - | **Moderate Drought** |
| **Chinook** | 6.45E-05 | 5.10E-03 | - | - |  |
| **Comet** | 1.68E-04 | 0.0195595 | 0.9061582 | - |  |
| **CTZ** | 1.37E-04 | 0.0222116 | 0.7518333 | 0.9975417 |  |
| **Centennial** | 0.090202 | - | - | - | **Severe Drought** |
| **Chinook** | 8.93E-03 | 0.8233731 | - | - |  |
| **Comet** | 4.19E-03 | 0.5369137 | 0.9794279 | - |  |
| **CTZ** | 0.9432712 | 0.3341307 | 0.0468495 | 0.0212125 |  |

### Linear Mixed Model and Weekly Growth Rates

ANOVA and Tukey post hoc tests revealed that treatment type (df= 2, F= 10.389, p= 3.18E-05), cultivar (df=4, F= 10.5697, p= 1.66E-08), and week (df= 10, F= 67.417, p <2.2E-16) significantly impacted weekly growth rates over the 12-week growing period (Table 5). Post-hoc testing exploring weeks as the fixed effect revealed that Weeks 2-4 (mid to late July) experienced significantly higher weekly growth rates compared to all other weeks during the 12-week growing period (Table 6). Post-hoc testing using treatment type as the fixed effect revealed that weekly growth rates did not significantly differ between the control and moderate treatment types (z= 0.363, p= 0.9298); however, significant differences were identified between control versus severe (z= 4.053, p= 0.0001) and moderate versus severe treatment types (z= 3.770, p= 0.0005). Tukey post-hoc tests exploring cultivar type as the fixed effect to evaluate which cultivars differed significantly in weekly growth rates revealed that while no cultivar types displayed significant differences in weekly growth rates compared to all other cultivars, CTZ was found to display significant differences with three out of four cultivars (Cascade, Chinook and Comet; Table 7).

**Table 5.** Summary statistics generated from a Type III ANOVA test using Satterthwaite’s method on outputs of the linear mixed model (LMM). Cultivar, treatment, and week served as fixed effects in the LMM, and weekly growth rates were response variables.

| **Fixed Effect** | **Sum Sq** | **Mean Sq** | **NumDF** | **DenDF** | **F-value** | **Pr(>F)** |
|---|---|---|---|---|---|---|
| **Cultivar** | 681.700 | 170.410 | 4 | 3,190.000 | 10.570 | 1.66E-08 |
| **Week** | 10,869.600 | 1,086.960 | 10 | 3,190.000 | 67.417 | <2.2E-16 |
| **Treatment** | 335.000 | 167.510 | 2 | 3,190.000 | 10.389 | 3.18E-05 |
| **Cultivar:Week** | 3,323.300 | 83.080 | 40 | 3,190.000 | 5.153 | <2.2E-16 |
| **Cultivar:Treatment** | 387.400 | 48.420 | 8 | 3,190.000 | 3.003 | 2.33E-03 |
| **Week:Treatment** | 1,773.600 | 88.680 | 20 | 3,190.000 | 5.500 | 3.75E-14 |
| **Cultivar:Week:Treatment** | 2,801.000 | 35.010 | 80 | 3,190.000 | 2.172 | 1.37E-08 |

**Table 6.** Matrix of p-values generated from Tukey’s post-hoc test using average growth rate values computed for each week over the 12-week growing period.

|  | **Week 2** | **Week 3** | **Week 4** | **Week 5** | **Week 6** | **Week 7** | **Week 9** | **Week 10** | **Week 11** | **Week 12** |
|---|---|---|---|---|---|---|---|---|---|---|
| **Week 1** | <0.0001 | <0.0001 | 0.0977 | 0.98 | 0.0357 | 0.0137 | <0.0001 | 0.9934 | 0.0241 | 0.9733 |
| **Week 2** | - | <0.0001 | <0.0001 | <0.0001 | <0.0001 | <0.0001 | <0.0001 | <0.0001 | <0.0001 | <0.0001 |
| **Week 3** | - | - | <0.0001 | <0.0001 | <0.0001 | <0.0001 | <0.0001 | <0.0001 | <0.0001 | <0.0001 |
| **Week 4** | - | - | - | 0.0013 | <0.0001 | <0.0001 | <0.0001 | 0.0026 | <0.0001 | 0.001 |
| **Week 5** | - | - | - | - | 0.5751 | 0.3756 | <0.0001 | 1.000 | 0.4884 | 1.000 |
| **Week 6** | - | - | - | - | - | 1.000 | 0.207 | 0.4562 | 1.000 | 0.6111 |
| **Week 7** | - | - | - | - | - | - | 0.3657 | 0.275 | 1.000 | 0.4089 |
| **Week 9** | - | - | - | - | - | - | - | <0.0001 | 0.2677 | 0.0001 |
| **Week 10** | - | - | - | - | - | - | - | - | 0.3744 | 1.000 |
| **Week 11** | - | - | - | - | - | - | - | - | - | 0.5242 |

**Table 7.** p-values generated from Tukey post hoc comparisons of the five tested cultivar types and their weekly growth rates over the 12-week growing period.

|  | **Centennial** | **Chinook** | **Comet** | **CTZ** |
|---|---|---|---|---|
| **Cascade** | 0.0625 | 0.1109 | 0.9931 | 0.0004 |
| **Centennial** | - | <0.0001 | 0.3073 | 0.4852 |
| **Chinook** | - | - | 0.0662 | <0.0001 |
| **Comet** | - | - | - | 0.011 |

Further post-hoc testing accounting for a treatment∼cultivar interaction revealed that under the control treatment, weekly growth rate did not significantly differ between any of the cultivar types (Table 8). However, under the moderate drought treatment type, Comet and CTZ displayed significant differences in weekly growth rates compared to Cascade (∼Comet, p= 0.0305; ∼CTZ, p= 0.0051) and Chinook (∼Comet, p= 0.0023; ∼CTZ, p= 0.0004), with CTZ and Comet (z= - 5.165, p <0.0001) eventually displaying significant differences in weekly growth rates under the severe drought regime (Table 8).

**Table 8.** p-values generated from Tukey post-hoc comparisons evaluating a treatment∼cultivar interaction using weekly growth rates as the response variable for each of the five cultivar types (Cascade, Centennial, Chinook, Comet and CTZ) under each of the three drought treatment types (control, moderate and severe).

|  | **Cascade** | **Centennial** | **Chinook** | **Comet** |  |
|---|---|---|---|---|---|
| **Centennial** | 0.9798 | - | - | - | **Control Treatment** |
| **Chinook** | 0.8933 | 0.6432 | - | - |  |
| **Comet** | 0.9969 | 1 | 0.8031 | - |  |
| **CTZ** | 0.8335 | 0.9837 | 0.4146 | 0.9854 |  |
| **Centennial** | 0.112 | - | - | - | **Moderate Drought** |
| **Chinook** | 0.6887 | 0.0089 | - | - |  |
| **Comet** | 0.0305 | 0.9069 | 0.0023 | - |  |
| **CTZ** | 0.0051 | 0.7265 | 0.0004 | 0.9995 |  |
| **Centennial** | 0.5047 | - | - | - | **Severe Drought** |
| **Chinook** | 0.2729 | 0.0071 | - | - |  |
| **Comet** | 0.0867 | 0.0003 | 0.9997 | - |  |
| **CTZ** | 0.0779 | 0.8428 | 0.0003 | <0.0001 |  |

### Leaf Surface Area and Leaf Circularity

Treatment (df= 2, F= 6.038, p= 0.01398), cultivar (df= 4, F= 35.638, p=6.85E-07), and treatment∼cultivar (df= 8, F= 5.733, p= 0.00292) interaction significantly impacted leaf surface area (Table 9). Post hoc testing examining treatment type as a fixed effect revealed that leaf surface area significantly differed between the moderate versus severe treatment types (p= 0.01297) but showed insignificant differences between the control and moderate treatment (p= 0.06977) as well as the control and severe treatment (p= 0.67813) types (Table 10), with the severe treatment type displaying the greatest variability in leaf surface area (Figure 3). When cultivar type was assessed as the fixed effect, our Tukey post-hoc test revealed that CTZ and Comet were significantly larger in leaf surface area than Cascade (∼CTZ, p= 0.000052; Comet, p= 0.000001), Centennial (∼CTZ, p= 0.000258; ∼Comet, p= 0.000006), and Chinook (∼CTZ, p= 0.011335; ∼Comet, p= 0.094796) (Figure 3b; Table 10). Additionally, Cascade was shown to display the least leaf surface area variability, while Comet displayed both the largest leaf surface area as well as the greatest variability (Figure 3b). Post-hoc testing focusing on treatment∼cultivar interaction showed that Cascade, Centennial, and Chinook most significantly differed in leaf surface area from Comet (∼Cascade, p= 0.000009; ∼Centennial, p= 0.000049; ∼Chinook= 0.000056) and CTZ (∼Cascade, p= 0.001782; ∼Centennial, p= 0.015704; ∼Chinook= 0.018768) under the severe drought treatment, with p-values supporting no significant difference in leaf surface area under the control or moderate drought treatments.

**Table 9.**
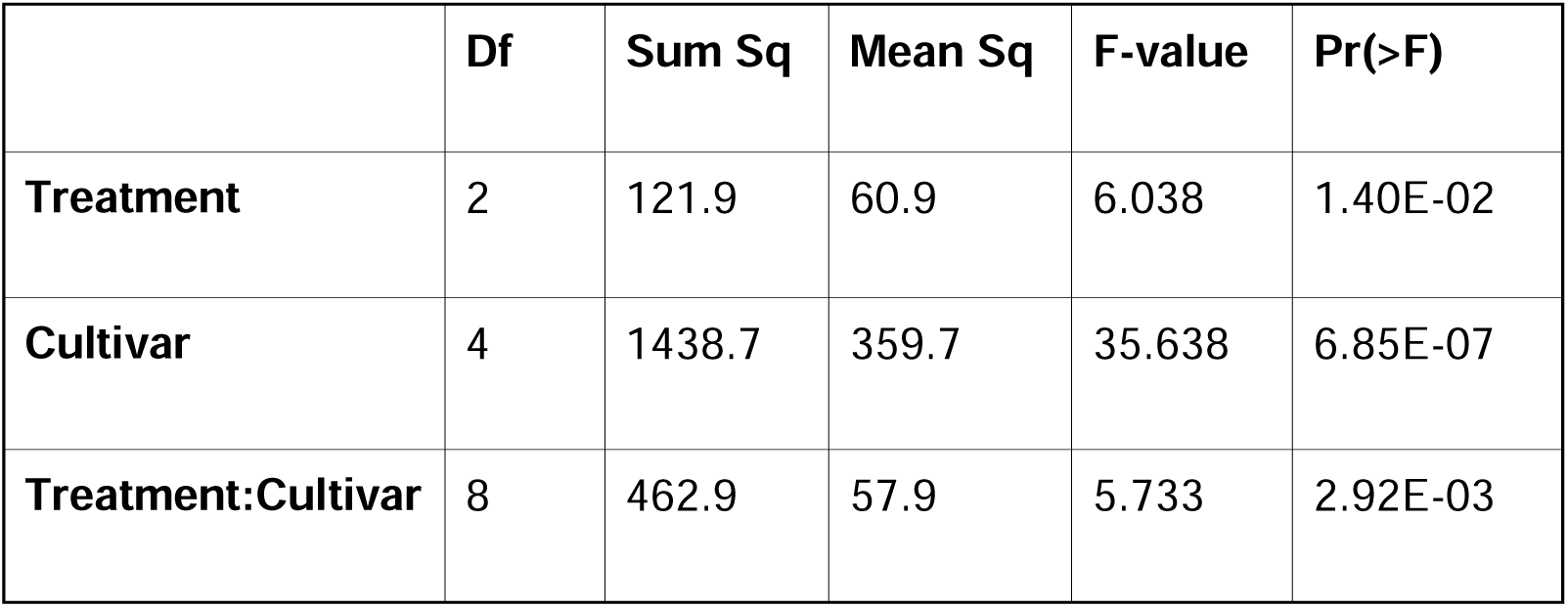
P-values generated from an ANOVA test using treatment, cultivar, and treatment:cultivar as fixed effects and surface area value as the response variable.

|  | **Df** | **Sum Sq** | **Mean Sq** | **F-value** | **Pr(>F)** |
|---|---|---|---|---|---|
| **Treatment** | 2 | 121.9 | 60.9 | 6.038 | 1.40E-02 |
| **Cultivar** | 4 | 1438.7 | 359.7 | 35.638 | 6.85E-07 |
| **Treatment:Cultivar** | 8 | 462.9 | 57.9 | 5.733 | 2.92E-03 |

**Figure 3.**
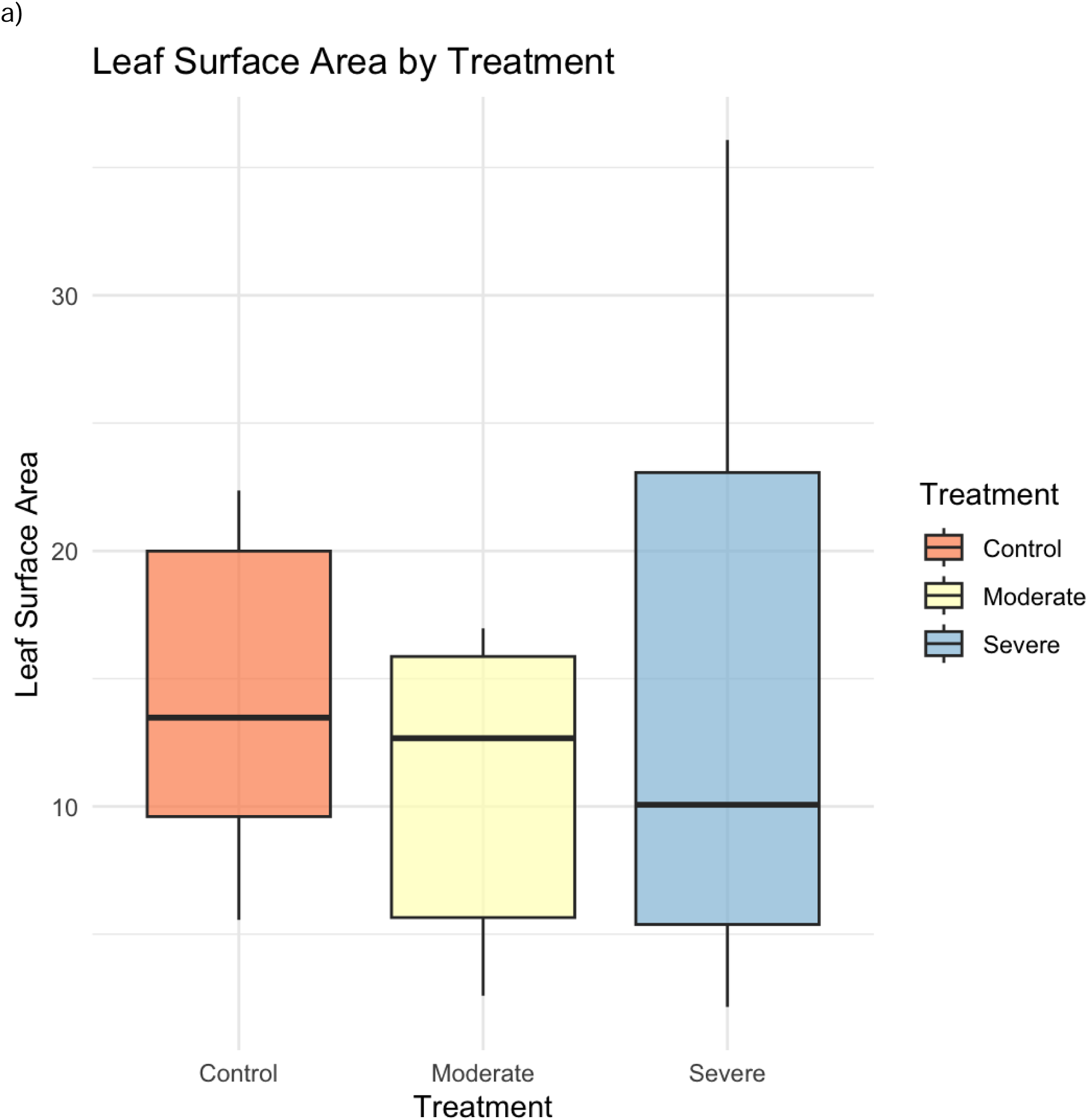
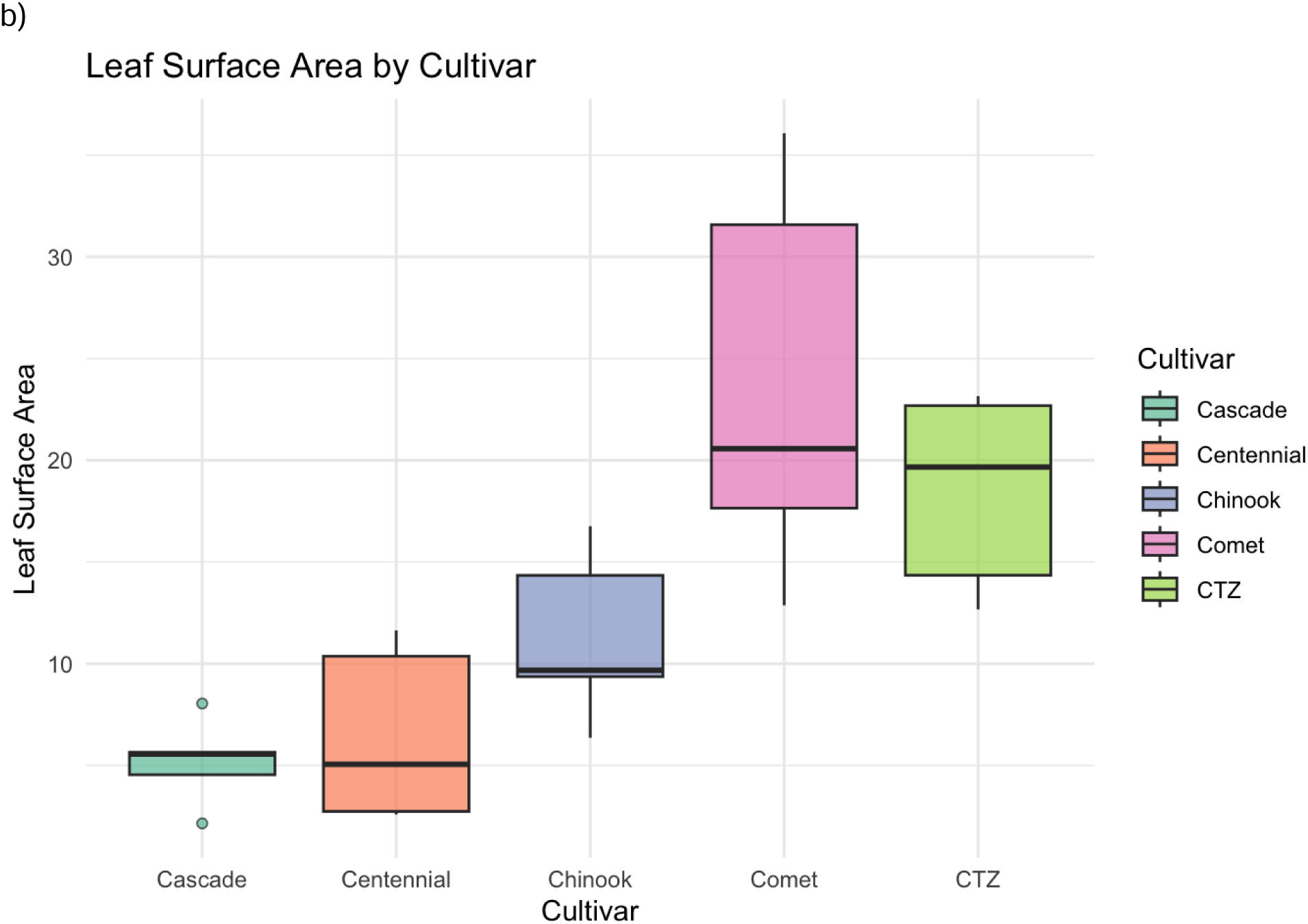
Boxplots of measured leaf surface area for each treatment type (a) or cultivar type (b).

**Table 10.** Adjusted p-value and summary statistics for Tukey’s post-hoc test using cultivar and treatment types as the fixed effects and leaf surface area as the response variable. *diff represents the* difference between the mean surface area values between groups, while *lwr* and *upr* values represent the lower and upper endpoints of the 95% confidence interval.

|  | Interaction | diff | lwr | upr | p.adj |
|---|---|---|---|---|---|
| **Treatment** | Control~Moderate | -3.676 | -7.630 | 0.279 | 0.070 |
|  | Control~Severe | 1.245 | -2.609 | 5.099 | 0.678 |
|  | Moderate~Severe | 4.920 | 1.066 | 8.774 | 0.013 |
| **Cultivar** | Cascade~Centennial | 2.024 | -4.302 | 8.351 | 0.847 |
|  | Cascade~Chinook | 6.656 | 0.599 | 12.713 | 0.029 |
|  | Cascade~Comet | 19.051 | 12.994 | 25.108 | 1.70E-06 |
|  | Cascade~CTZ | 13.955 | 7.898 | 20.012 | 5.29E-05 |
|  | Centennial~Chinook | 4.632 | -1.425 | 10.689 | 0.174 |
|  | Centennial~Comet | 17.027 | 10.970 | 23.084 | 6.10E-06 |
|  | Centennial~CTZ | 11.931 | 5.874 | 17.988 | 2.59E-04 |
|  | Chinook~Comet | 12.395 | 6.620 | 18.170 | 1.10E-04 |
|  | Chinook~CTZ | 7.298 | 1.523 | 13.074 | 0.011 |
|  | Comet~CTZ | -5.097 | -10.872 | 0.679 | 0.095 |

ANOVA test of leaf circularity revealed that while treatment type (df= 2, F= 2.053, p= 0.16797) and a treatment∼cultivar (df= 8, F= 1.552, p= 0.23094) did not significantly impact leaf circularity, cultivar type (df= 4, F= 2.053, p= 2.47E-03) was found to have a significant impact on leaf circularity (Figure 4; Table 11). Post hoc testing of cultivar type revealed that Comet displayed significantly lower leaf circularity compared to Cascade (p= 0.010855), Centennial (p= 0.004089), and Chinook (p= 0.009506) and was insignificantly lower when compared to CTZ (p= 0.273186) (Figure 4b; Table 12).

**Figure 4.**
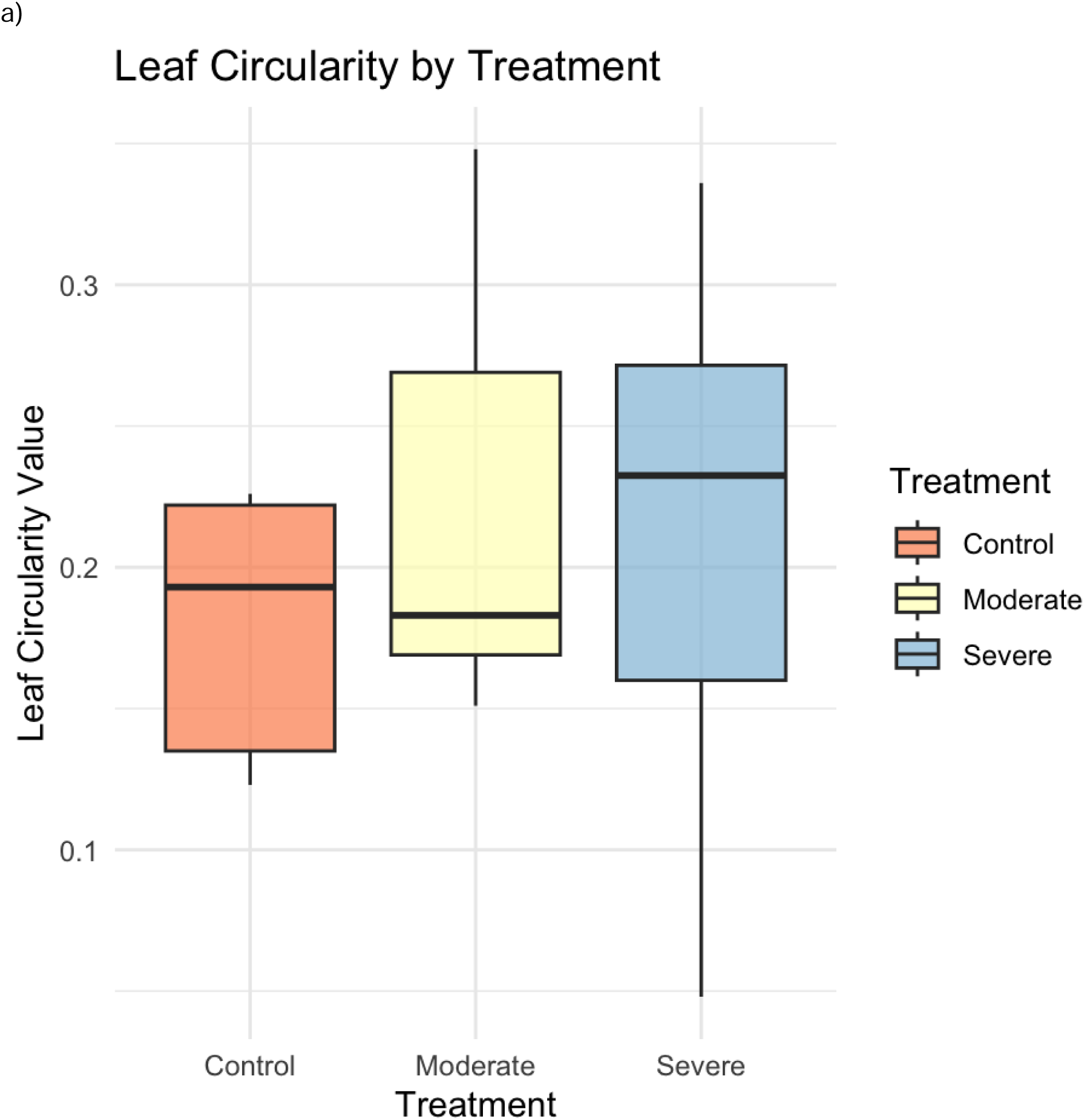
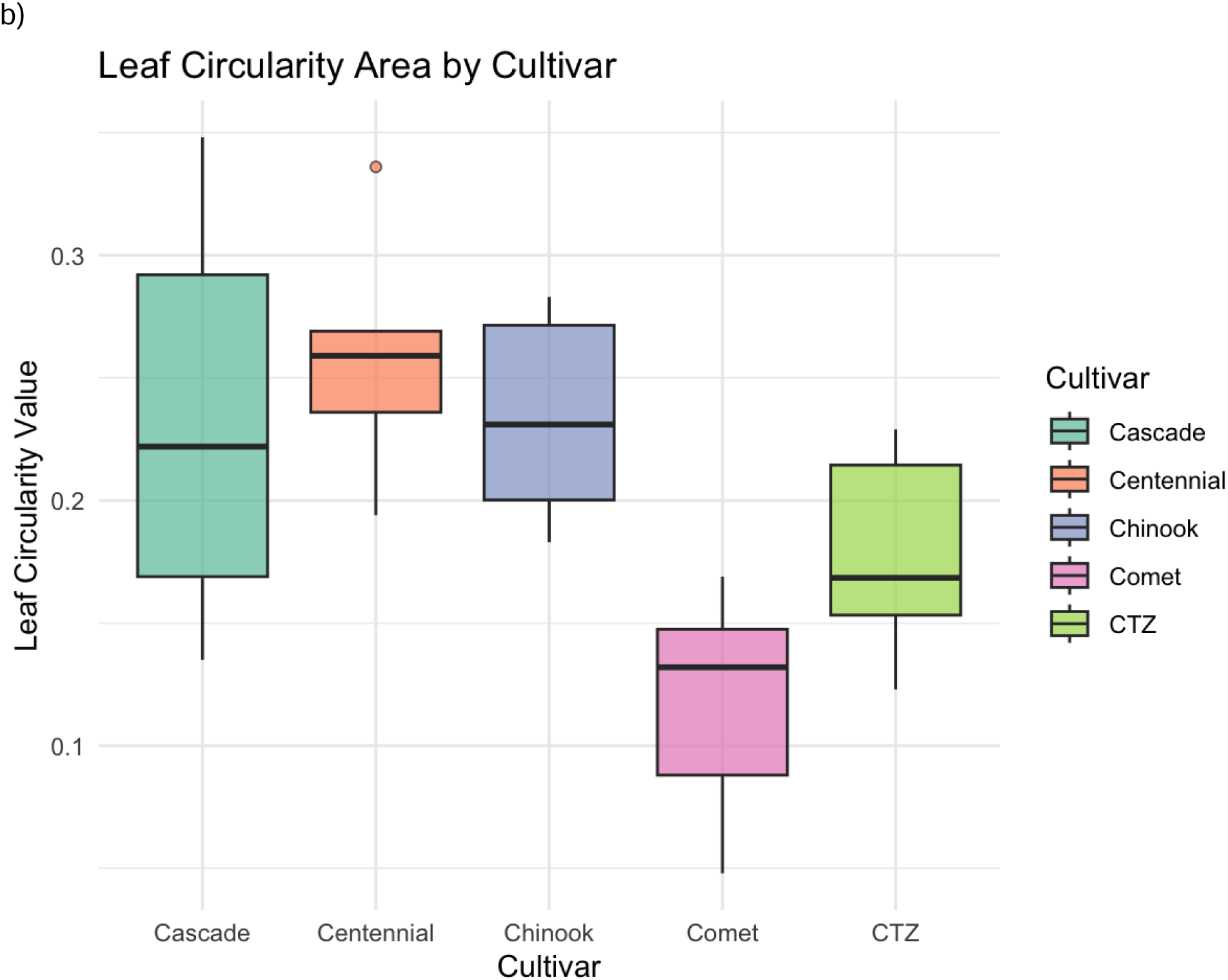
Box plots of leaf circularity by treatment type (a) or cultivar type (b). Leaf circularity values were calculated using the following formula: (4π ×Area)/Perimeter^2, as suggested by Hightower et al. 2024.

**Table 11.** Summary statistics and p-values from an ANOVA test using leaf circularity as the response variable and treatment, cultivar, and treatment:cultivar as the fixed effects.

|  | **Df** | **Sum Sq** | **Mean Sq** | **F-value** | **Pr(>F)** |
|---|---|---|---|---|---|
| **Treatment** | 2 | 0.00997 | 0.004985 | 2.053 | 0.168 |
| **Cultivar** | 4 | 0.07179 | 0.017947 | 7.391 | 2.47E-03 |
| **Treatment:Cultivar** | 8 | 0.03016 | 0.00377 | 1.552 | 0.231 |

**Table 12.** Summary statistics and adjusted p-values generated from a Tukey post-hoc test exploring treatment type and cultivar as fixed effects and leaf circularity scores as the response variable.

|  | Interaction | diff | lwr | upr | p.adj |
|---|---|---|---|---|---|
| **Treatment** | Control~Moderate | 0.046 | -0.015 | 0.107 | 0.155 |
|  | Control~Severe | 0.031 | -0.029 | 0.091 | 0.385 |
|  | Moderate~Severe | -0.015 | -0.075 | 0.045 | 0.785 |
| **Cultivar** | Cascade~Centennial | 0.016 | -0.082 | 0.115 | 0.983 |
|  | Cascade~Chinook | -0.003 | -0.097 | 0.091 | 1.000 |
|  | Cascade~Comet | -0.119 | -0.213 | -0.025 | 1.09E-02 |
|  | Cascade~CTZ | -0.059 | -0.153 | 0.035 | 0.322 |
|  | Centennial~Chinook | -0.020 | -0.114 | 0.074 | 0.961 |
|  | Centennial~Comet | -0.136 | -0.230 | -0.042 | 4.09E-03 |
|  | Centennial~CTZ | -0.076 | -0.170 | 0.018 | 0.141 |
|  | Chinook~Comet | -0.116 | -0.206 | -0.026 | 9.51E-03 |
|  | Chinook~CTZ | -0.056 | -0.146 | 0.034 | 0.332 |
|  | Comet~CTZ | 0.060 | -0.030 | 0.150 | 0.273 |

### Soil Microbial and Fungal Community Dynamics in Response to Drought

From ITS sequencing, we were able to recover 97 fungal OTUs found across our 28 sampled hops rhizospheres. ANOVA testing using percent abundance data generated from ITS sequencing found that neither cultivar (df= 4, F= 0.0079, p= 0.9999), treatment (df= 2, F= 0.0038, p= 0.9962), nor timepoint (df= 1, F= 0.0165, p= 0.8979) impacted fungal composition within the rhizosphere (Table 13). However, accounting for an interaction between percent abundance for OTU and each of our fixed effects, ANOVA testing revealed that OTU percent abundance did identified did significantly differ between cultivar type (df= 384, F= 2.418, p< 2E-16) and timepoint (df= 384, F= 2.7637, p= 1.03E-11) but not treatment type (df= 384, F= 0.6369, p= 1.0000) (Table 13). Post hoc testing accounting for a cultivar∼OTU_identified interaction revealed that three fungal families, Aspergillaceae (t= 6.575, p= 5.54E-11), Cephalothecaceae (t= 4.154, p= 3.33E-05), and Cladosporiaceae (t= 16.366, p< 2E-16) displayed significant differences in percent abundance between cultivar types.

**Table 13.**
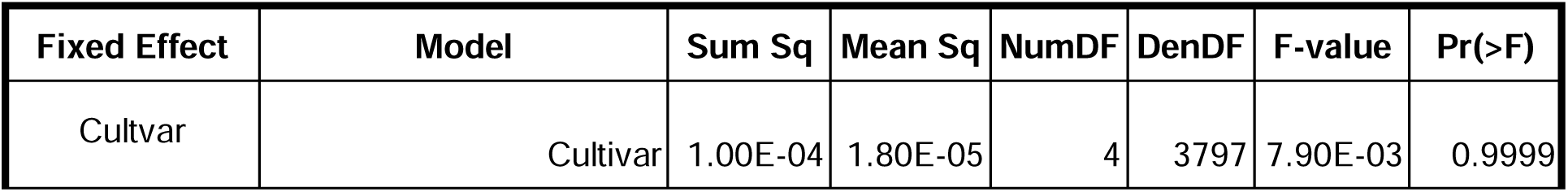
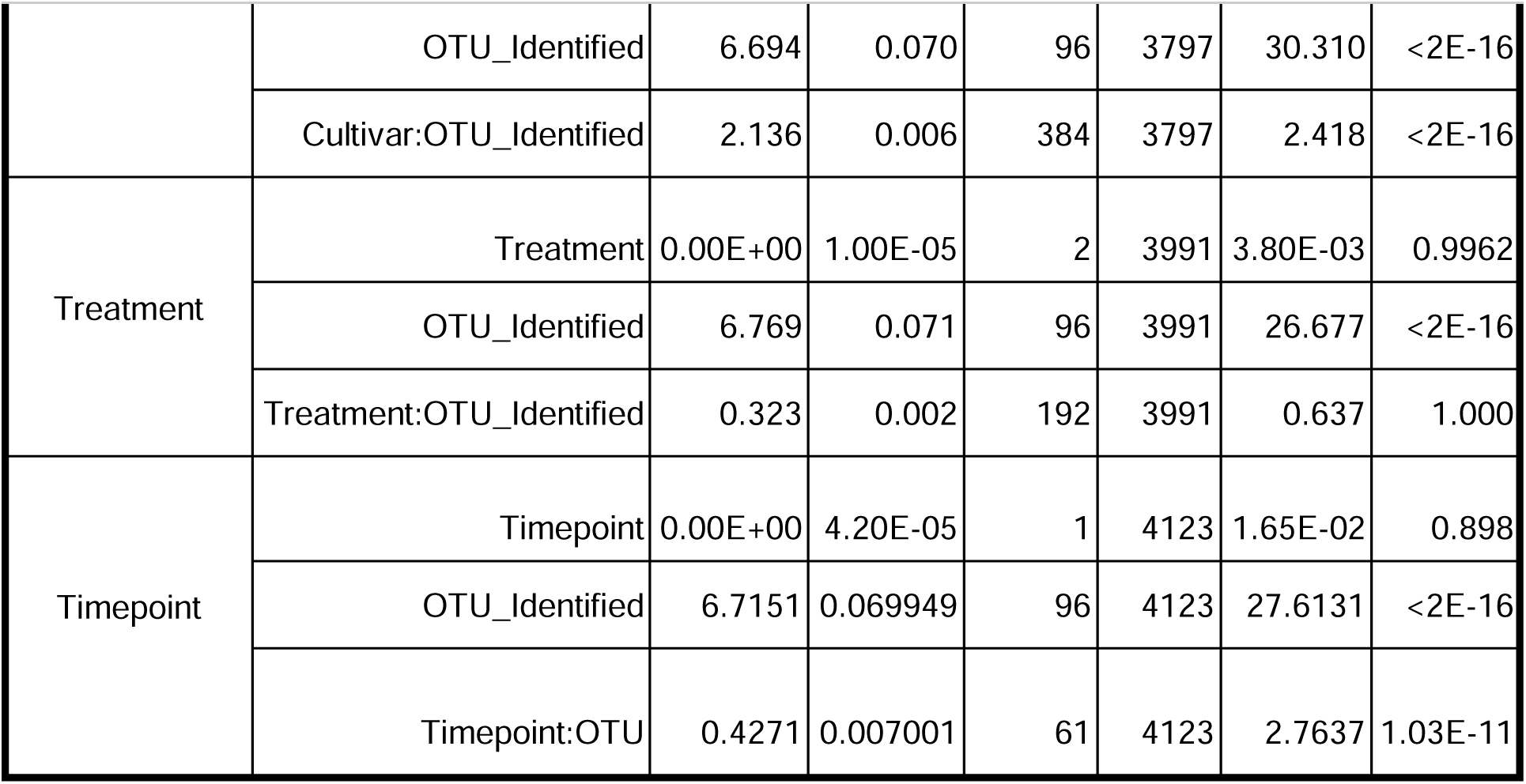
Summary statistics and p-values generated from an ANOVA test of fungal OTU composition identified from collected soil samples via ITS sequencing. Cultivar, treatment type, and time point (start vs end) were treated as the fixed effects, while composition of identified fungal OTUs was the response variable.

| Fixed Effect | Model | Sum Sq | Mean Sq | NumDF | DenDF | F-value | Pr(>F) |
|---|---|---|---|---|---|---|---|
| Cultvar | Cultivar | 1.00E-04 | 1.80E-05 | 4 | 3797 | 7.90E-03 | 0.9999 |
|  | OTU_Identified | 6.694 | 0.070 | 96 | 3797 | 30.310 | <2E-16 |
|  | Cultivar:OTU_Identified | 2.136 | 0.006 | 384 | 3797 | 2.418 | <2E-16 |
| Treatment | Treatment | 0.00E+00 | 1.00E-05 | 2 | 3991 | 3.80E-03 | 0.9962 |
|  | OTU_Identified | 6.769 | 0.071 | 96 | 3991 | 26.677 | <2E-16 |
|  | Treatment:OTU_Identified | 0.323 | 0.002 | 192 | 3991 | 0.637 | 1.000 |
| Timepoint | Timepoint | 0.00E+00 | 4.20E-05 | 1 | 4123 | 1.65E-02 | 0.898 |
|  | OTU_Identified | 6.7151 | 0.069949 | 96 | 4123 | 27.6131 | <2E-16 |
|  | Timepoint:OTU | 0.4271 | 0.007001 | 61 | 4123 | 2.7637 | 1.03E-11 |

**Table 14.** Summary statistics and p-values generated from ANOVA test of archaeal OTU composition identified from collected soil samples via 16Ssequencing. Cultivar, treatment type, and time point (start vs end) were treated as the fixed effects, while composition of identified fungal OTU was the response variable.

| Fixed Effect | Model | Sum Sq | Mean Sq | NumDF | DenDF | F-value | Pr(>F) |
|---|---|---|---|---|---|---|---|
| Cultvar | Cultivar | 4.33E-06 | 1.08E-06 | 4 | 306 | 1.94 | 0.1039826 |
|  | OTU_Identified | 4.04E-05 | 8.09E-06 | 5 | 306 | 14.49 | 9.63E-13 |
|  | Cultivar:OTU_Identified | 3.03E-05 | 1.51E-06 | 20 | 306 | 2.71 | 1.41E-04 |
| Treatment | Treatment | 3.02E-07 | 1.51E-07 | 2 | 318 | 0.24 | 0.79 |
|  | OTU_Identified | 4.34E-05 | 8.69E-06 | 5 | 318 | 13.55 | 5.37E-12 |
|  | Treatment:OTU_Identified | 1.33E-06 | 1.33E-07 | 10 | 318 | 0.21 | 0.996 |
| Timepoint | Timepoint | 1.21E-06 | 1.21E-06 | 1 | 324 | 2.03 | 0.155 |
|  | OTU_Identified | 4.41E-05 | 8.82E-06 | 5 | 324 | 14.86 | 3.95E-13 |
|  | Timepoint:OTU | 1.18E-05 | 2.37E-06 | 5 | 324 | 3.98 | 1.61E-03 |

Aspergillaceae (t= 18.303, p< 2E-16), Cephalothecaceae (t= 7.572, p= 4.50E-14), and Cladosporiaceae (t= 21.069, p< 2E-16) were identified as statistically impacted in percent abundance between time points by our LMM, as well as the addition of Chaetomiaceae (t= 3.219, p= 1.30E-03) and Microascaceae (t= 3.597, p= 3.26E-04).

Using targeted 16S sequencing, we recovered 6 archaeal OTU’s across all sampled rhizospheres. Neither cultivar (df= 4, F= 1.9382, p= 0.1039826), treatment type (df= 2, F= 0.2358, p= 0.7900), nor time point (df= 1, F= 2.0341, p= 0.154767) independently impacted archaeal percent abundance, however, did affect the composition of specific archaeal families (Table 13). Post hoc testing revealed that of the six identified archaeal OTUs, only one clade, Thaumarchaeota, was found to significantly differ in percent abundance across cultivar types and between timepoints. Using timepoint as our fixed factor, Thaumarchaeota percent abundance was significantly higher (t= 3.269, p= 0.0012) at the start of the experiment than at the end of the twelve weeks (Fig. 5). Additionally, post-hoc testing using cultivar as the fixed effect unveiled that two cultivars, Chinook (t= 3.005, p= 0.00288) and Comet (t= 4.402, p= 1.48E-05), displayed significantly higher percent abundance of Thaumarchaeota compared to the other three tested cultivar types (Fig 5).

**Figure 5.**
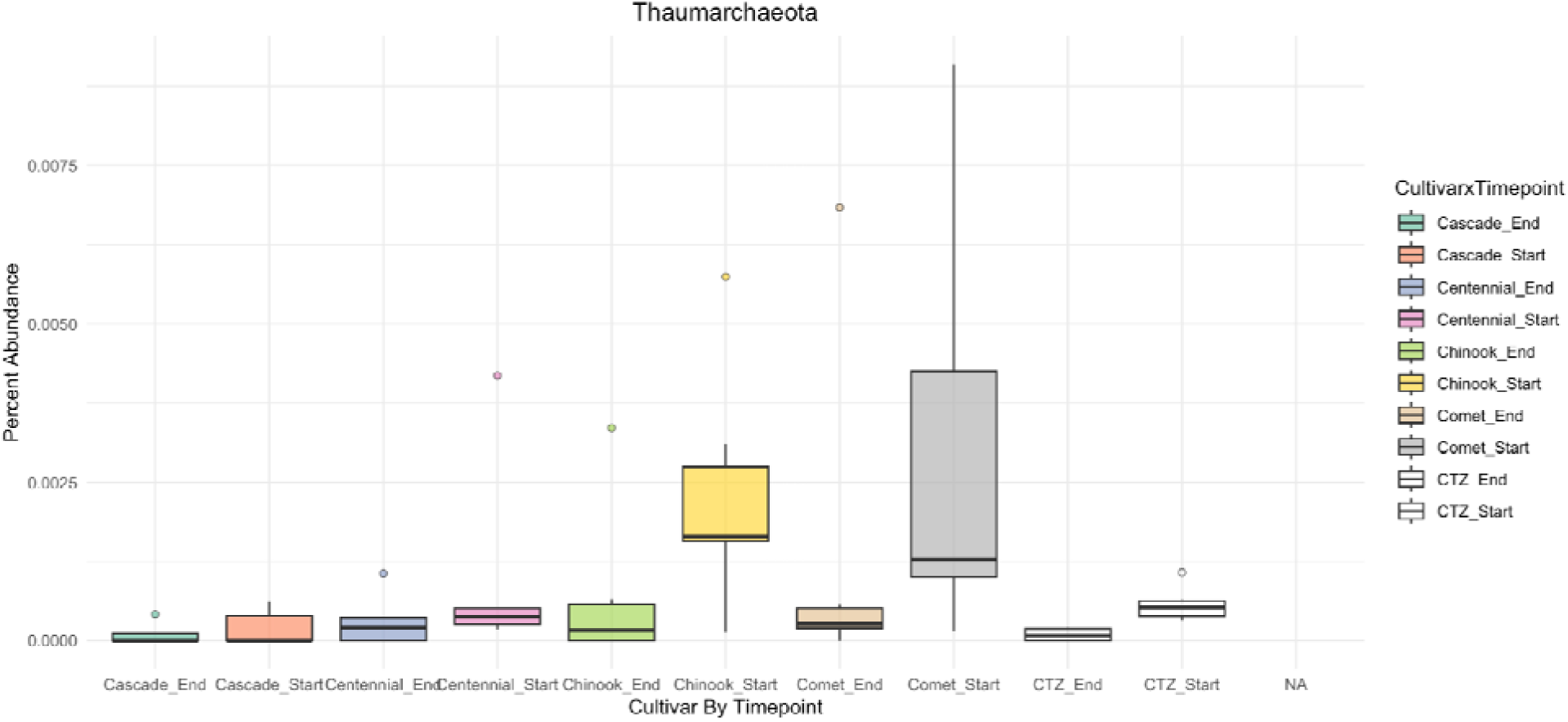
Boxplot representing the average percent abundance of the archaeal phylum Thaumarchaeota across the five test cultivar types (Cascade, Centennial, Chinook, Comet, and CTZ) between two time points, the start and end of the 12-week drought experimental period.

16S sequencing targeting bacteria recovered 175 bacterial OTUs across our sampled hops rhizospheres. ANOVA tests using cultivar (df= 4, F= 0.2684, p= 0.8984), treatment type (df= 2, F= 0.1436, p= 0.8662), and timepoint (df= 2, F= 0.1639, p= 0.6856) revealed that, independently, these three fixed effects did not significantly impact total bacterial OTU percent abundance; however, each effect was found to significantly impact particular OTUs’ percent abundance independently (Table 15). Using timepoint as a fixed factor, over half (64.5%; n=111) of the identified bacterial families showed a significant (p-value < 0.05) difference in their total percent abundance, with nine bacterial families displaying highly significant (p-value < 0.001) changes in percent abundance during the 12-week experimental period.

**Table 15.** Summary statistics and p-values generated from an ANOVA test of bacterial OTU composition identified from collected soil samples via 16S sequencing. Cultivar, treatment type, and time point (start vs end) were treated as the fixed effects, while composition of identified fungal OTUs was the response variable.

| Fixed Effect | Model | Sum Sq | Mean Sq | NumDF | DenDF | F-value | Pr(>F) |
|---|---|---|---|---|---|---|---|
| Cultvar | Cultivar | 1.30E-04 | 3.22E-05 | 4 | 8981 | 0.27 | 0.8984 |
|  | OTU_Identified | 2.66E+00 | 1.53E-02 | 174 | 8981 | 127.21 | <2E-16 |
|  | Cultivar:OTU_Identified | 3.17E-01 | 4.55E-04 | 696 | 8981 | 3.79 | <2E-16 |
| Treatment | Treatment | 4.00E-05 | 2.04E-05 | 2 | 9331 | 0.14 | 0.8662 |
|  | OTU_Identified | 2.68E+00 | 1.54E-02 | 174 | 9331 | 108.26 | <2E-16 |
|  | Treatment:OTU_Identified | 6.75E-02 | 1.94E-04 | 348 | 9331 | 1.36 | 1.17E-05 |
| Timepoint | Timepoint | 2.00E-05 | 2.33E-05 | 1 | 9506 | 0.16 | 0.686 |
|  | OTU_Identified | 2.67E+00 | 1.53E-02 | 174 | 9506 | 108.11 | <2E-16 |
|  | Timepoint:OTU | 4.54E-02 | 2.61E-04 | 174 | 9506 | 1.84 | 2.05E-10 |

Using cultivar as a fixed effect, seven bacterial OTU’s significantly differed in percent abundance between cultivar types; Bacillaceae (t= 37.873, p< 2E-16), Nocardioidaceae (t= 4.847, p=1.28E-06), Paenibacillaceae (t= 21.181, p< 2E-16), Planctomycetaceae (t= 2.008, p= 4.47E-02), Rhodothermaceae (t= -2.101, p= 3.57E-02), Streptomycetaceae (t= 4.967, p= 6.92E-07) and Streptosporangiaceae (t= 8.633, p< 2E-16). Of the seven bacterial OTU’s identified as varying significantly between cultivar types, only three OTU’s, Bacillaceae, Paenibacillaceae, and Streptosporangiaceae, showed significant differences in percent abundance for each cultivar type, whereas other bacterial OTUs such as Planctomycetaceae displayed significant differences in percent abundance for only one particular cultivar, possibly displaying novel associations (Table 16; Figure 6).

**Table 16.** Summary statistics and p-values associated with each cultivar type for bacterial OTU’s identified as significant by ANOVA testing.

| **Cultivar** | **OTU** | **Estimate** | **Std Error** | **df** | **t-value** | **Pr(>\|t\|)** |
|---|---|---|---|---|---|---|
| **Centennial** | Bacillaceae | -1.37E-02 | 6.93E-03 | 8981 | -1.982 | 4.76E-02 |
|  | Nocardioidaceae | -1.08E-03 | 6.93E-03 | 8981 | -0.156 | 8.76E-01 |
|  | Paenibacillaceae | -2.87E-02 | 6.93E-03 | 8981 | -4.14 | 3.51E-05 |
|  | Planctomycetaceae | 1.06E-03 | 6.93E-03 | 8981 | 0.153 | 0.878298 |
|  | Rhodothermaceae | -1.16E-03 | 6.00E-03 | 8981 | -0.192 | 8.47E-01 |
|  | Streptomycetaceae | 1.30E-02 | 6.93E-03 | 8981 | 1.882 | 5.99E-02 |
|  | Streptosporangiaceae | -1.32E-03 | 6.93E-03 | 8981 | -0.19 | 8.49E-01 |
| **Chinook** | Bacillaceae | -8.70E-02 | 6.64E-03 | 8981 | -13.109 | 2.00E-16 |
|  | Nocardioidaceae | -8.49E-03 | 6.64E-03 | 8981 | -1.279 | 2.01E-01 |
|  | Paenibacillaceae | -3.61E-02 | 6.64E-03 | 8981 | -5.435 | 5.64E-08 |
|  | Planctomycetaceae | 1.35E-02 | 6.64E-03 | 8981 | 2.038 | 4.16E-02 |
|  | Rhodothermaceae | 2.41E-03 | 5.75E-03 | 8981 | 0.42 | 6.74E-01 |
|  | Streptomycetaceae | 2.51E-02 | 6.64E-03 | 8981 | 3.789 | 1.53E-04 |
|  | Streptosporangiaceae | -1.80E-02 | 6.64E-03 | 8981 | -2.705 | 6.84E-03 |
| **Comet** | Bacillaceae | -2.05E-02 | 6.64E-03 | 8981 | -3.096 | 1.97E-03 |
|  | Nocardioidaceae | 9.13E-03 | 6.64E-03 | 8981 | 1.376 | 1.69E-01 |
|  | Paenibacillaceae | -3.97E-02 | 6.64E-03 | 8981 | -5.988 | 2.21E-09 |
|  | Planctomycetaceae | 9.25E-03 | 6.64E-03 | 8981 | 1.393 | 1.64E-01 |
|  | Rhodothermaceae | 9.89E-04 | 5.75E-03 | 8981 | 0.172 | 8.63E-01 |
|  | Streptomycetaceae | 4.73E-03 | 6.64E-03 | 8981 | 0.712 | 4.76E-01 |
|  | Streptosporangiaceae | -2.14E-02 | 6.64E-03 | 8981 | -3.225 | 1.26E-03 |
| **CTZ** | Bacillaceae | 1.13E-01 | 6.64E-03 | 8981 | 17.053 | 2.00E-16 |
|  | Nocardioidaceae | 1.10E-02 | 6.64E-03 | 8981 | 1.653 | 9.85E-02 |
|  | Paenibacillaceae | -4.44E-02 | 6.64E-03 | 8981 | -6.693 | 2.32E-11 |
|  | Planctomycetaceae | 4.95E-03 | 6.64E-03 | 8981 | 0.746 | 4.56E-01 |
|  | Rhodothermaceae | 5.14E-03 | 5.75E-03 | 8981 | 0.894 | 3.72E-01 |
|  | Streptomycetaceae | 8.24E-03 | 6.64E-03 | 8981 | 1.242 | 2.14E-01 |
|  | Streptosporangiaceae | -3.04E-02 | 6.64E-03 | 8981 | -4.584 | 4.61E-06 |

**Figure 6.**
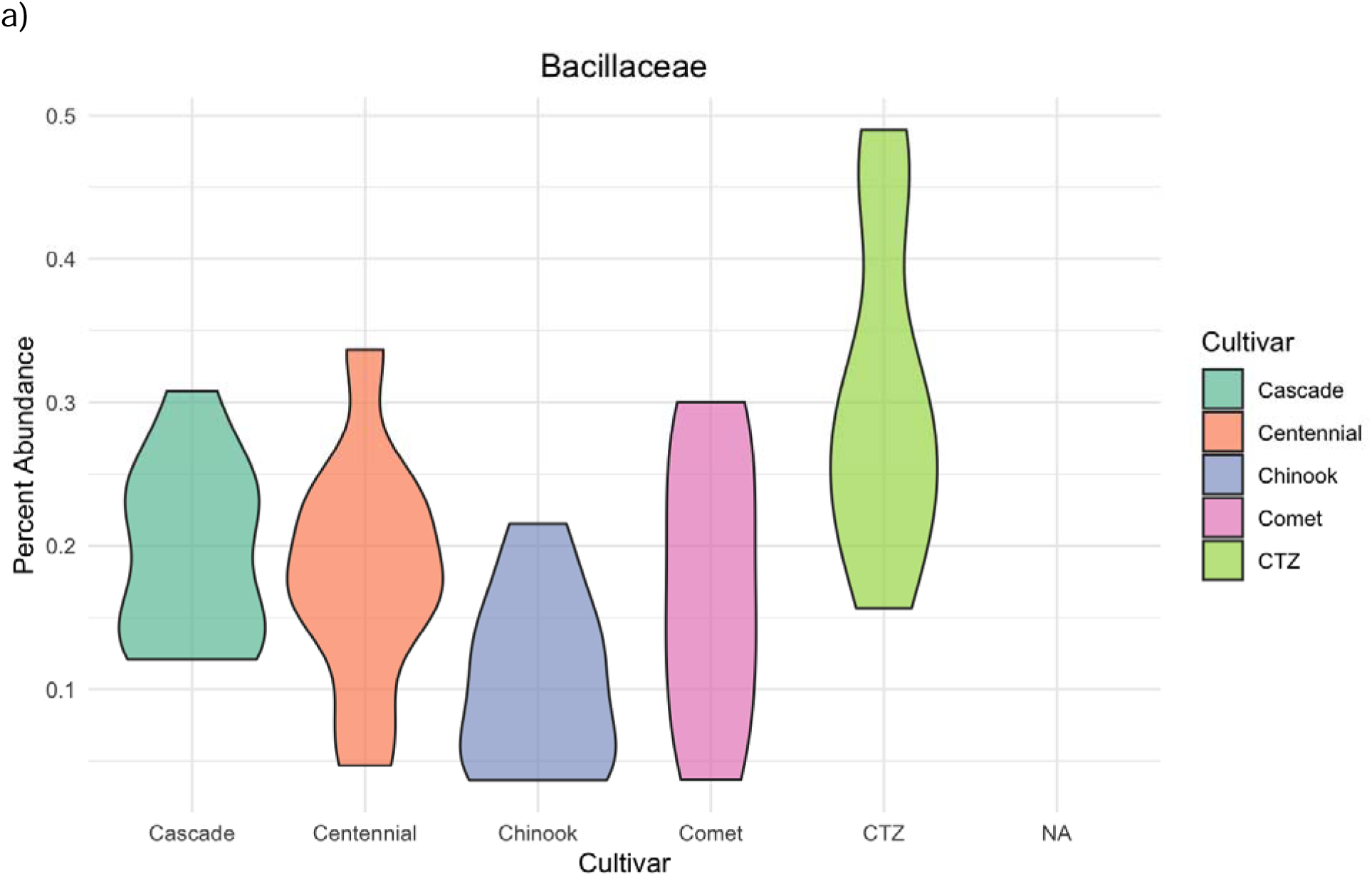
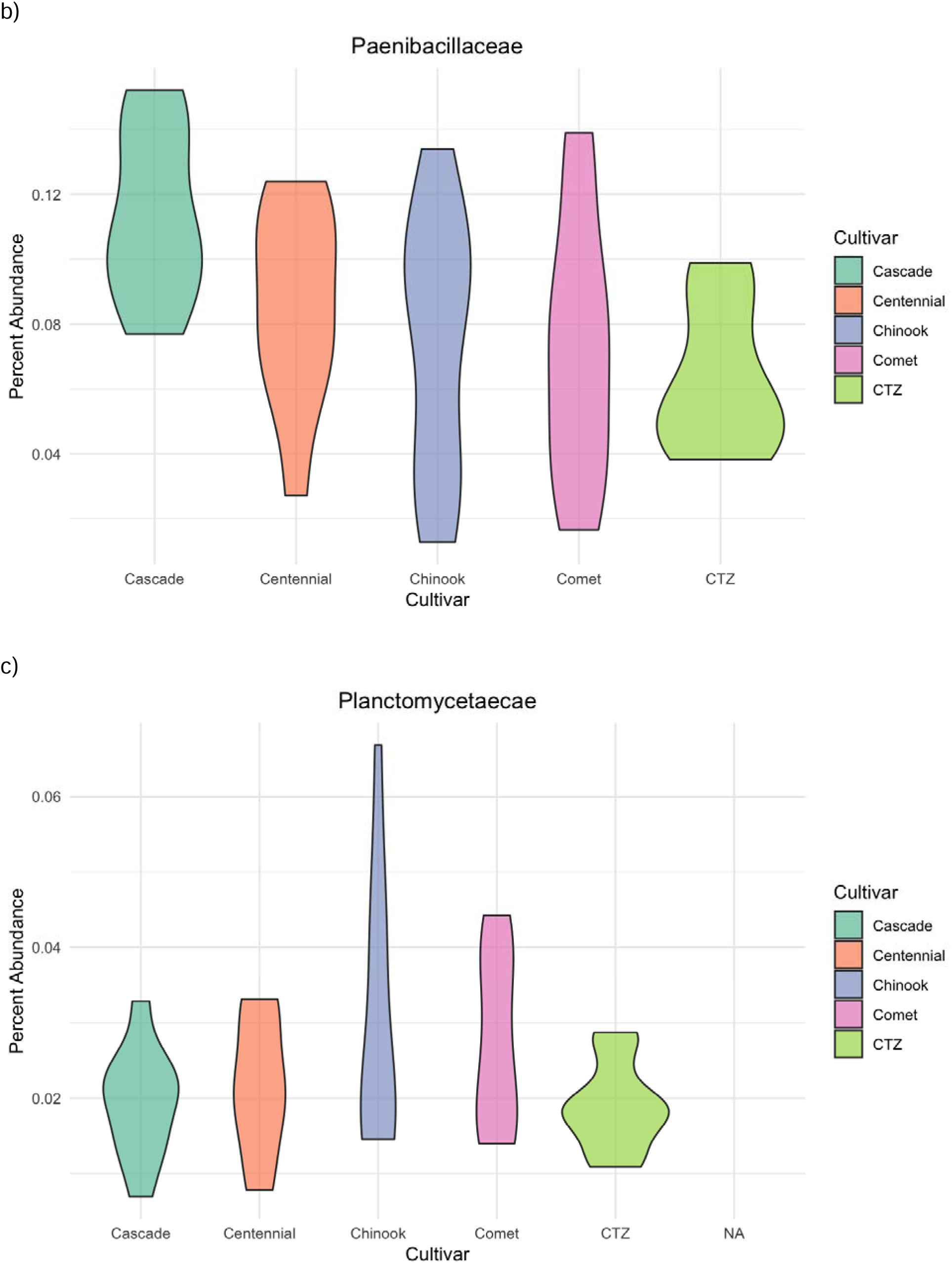
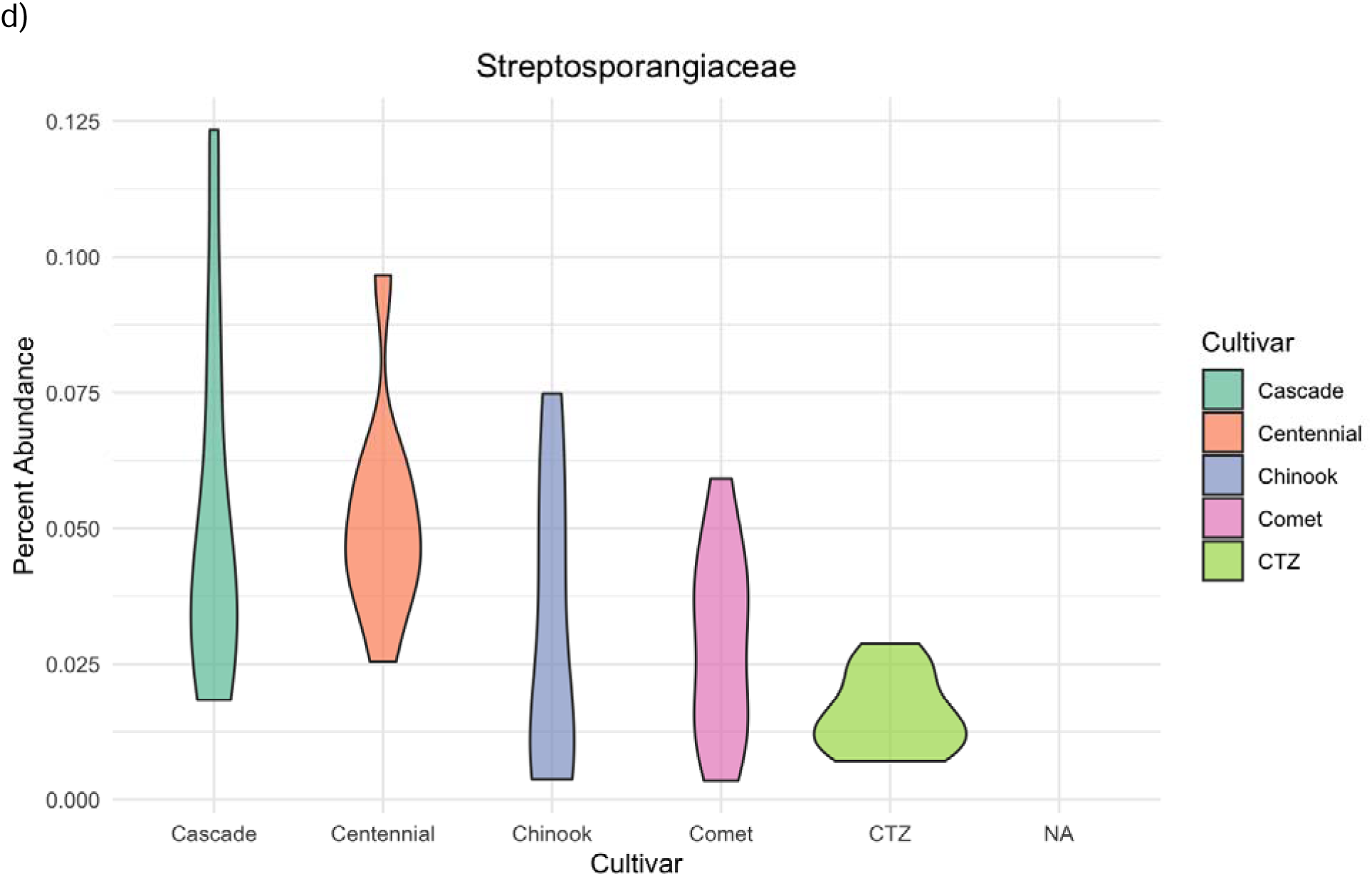
Violin plots of hops cultivars’ percent abundance across the five cultivar types for the following bacterial families: a) Bacillaceae, b) Paenibacillaceae, c) Planctomycetaceae, d) Streptosporangiacae

Using treatment as the fixed effect, eight bacterial OTU’s significantly differed in percent abundance between the start and end of the experiment (Table 17), several of which have been detected between other fixed factors as well, as described above. When the LMM was examined for treatment∼OTU interaction as the fixed effect, only one bacterial OTU significantly differed in percent abundance, Bacillaceae (moderate treatment, t= 3.191, p= 0.00142; severe treatment, t= -4.342, p= 1.42E-05). Visualization of percent abundance between treatment types revealed that for Bacillaceae, percent abundance was highest in the moderate drought regime and lowest in the severe regime, with the control treatment having a significantly lower percent abundance compared to the moderate treatment (Figure 7).

**Table 17.**
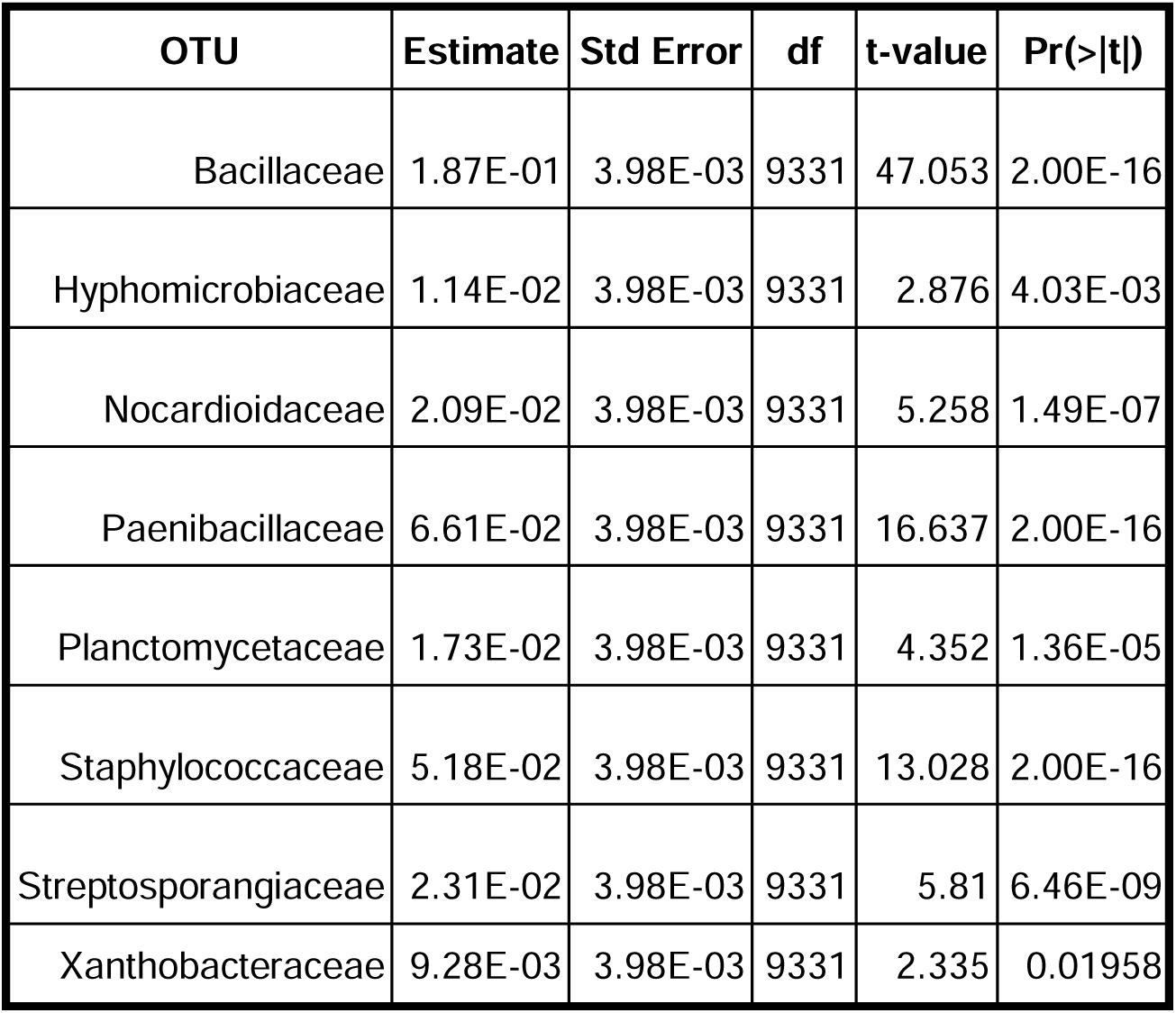
Table of eight bacterial OTU’s detected from the rhizosphere and identified as displaying a significant change in percent abundance between the start and end of the 12-week growing period. The table displays summary statistics and associated p-values generated from a linear mixed model using treatment and OTU as the fixed effects and percent abundance as the response variable.

| **OTU** | **Estimate** | **Std Error** | **df** | **t-value** | **Pr(>\|t\|)** |
|---|---|---|---|---|---|
| Bacillaceae | 1.87E-01 | 3.98E-03 | 9331 | 47.053 | 2.00E-16 |
| Hyphomicrobiaceae | 1.14E-02 | 3.98E-03 | 9331 | 2.876 | 4.03E-03 |
| Nocardioidaceae | 2.09E-02 | 3.98E-03 | 9331 | 5.258 | 1.49E-07 |
| Paenibacillaceae | 6.61E-02 | 3.98E-03 | 9331 | 16.637 | 2.00E-16 |
| Planctomycetaceae | 1.73E-02 | 3.98E-03 | 9331 | 4.352 | 1.36E-05 |
| Staphylococcaceae | 5.18E-02 | 3.98E-03 | 9331 | 13.028 | 2.00E-16 |
| Streptosporangiaceae | 2.31E-02 | 3.98E-03 | 9331 | 5.81 | 6.46E-09 |
| Xanthobacteraceae | 9.28E-03 | 3.98E-03 | 9331 | 2.335 | 0.01958 |

**Figure 7.**
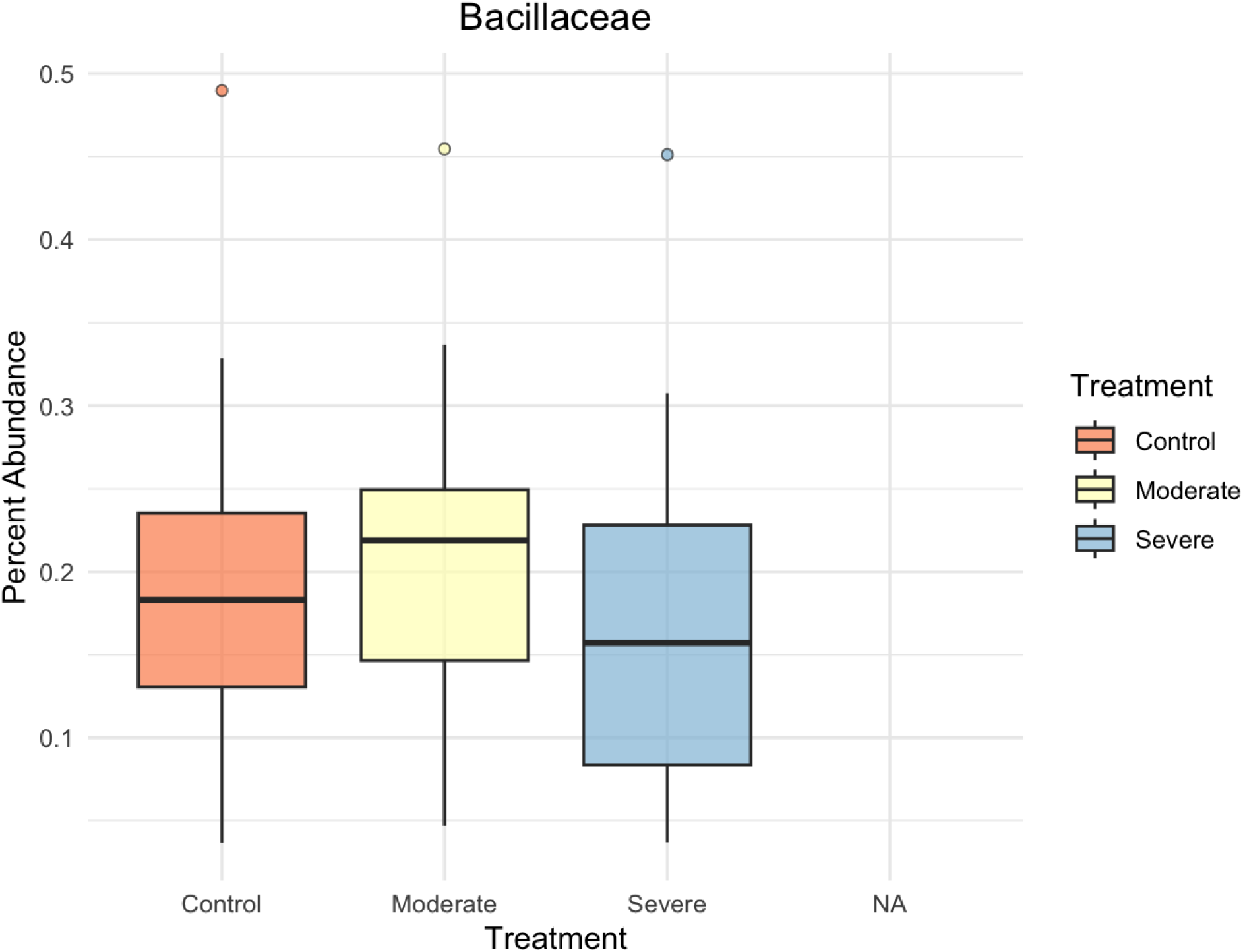
Boxplot representing the bacterial family Bacillaceae measured percent abundance for each of the following treatment types: control, moderate, and severe.

### Impact on Rhizospheric Nutrient Bioavailability

ANOVA testing of five major bioavailable nutrients (PO4, NH4, NO3, DIN and DOC) revealed cultivar type significantly impacted nutrient bioavailability for four of the five nutrient types. PO4 (df= 4, F= 3.159, p= 0.045), NO3 (df= 4. F= 14.211, p< 0.001), DIN (df= 4, F= 7.136, p= 0.002) and DOC (df= 4, F= 5.576, p= 0.006) bioavailability were found to be significantly impacted by cultivar type. Under both control conditions and the moderate drought regime, CTZ displayed the greatest amount of extractable PO4 compared to other cultivar types (Figure 8a). However, under severe drought conditions, Comet not only displayed the greatest amount of bioavailable PO4 compared to other cultivar types, but displayed the highest overall amount when compared to moderate drought and control treatments.

**Figure 8.**
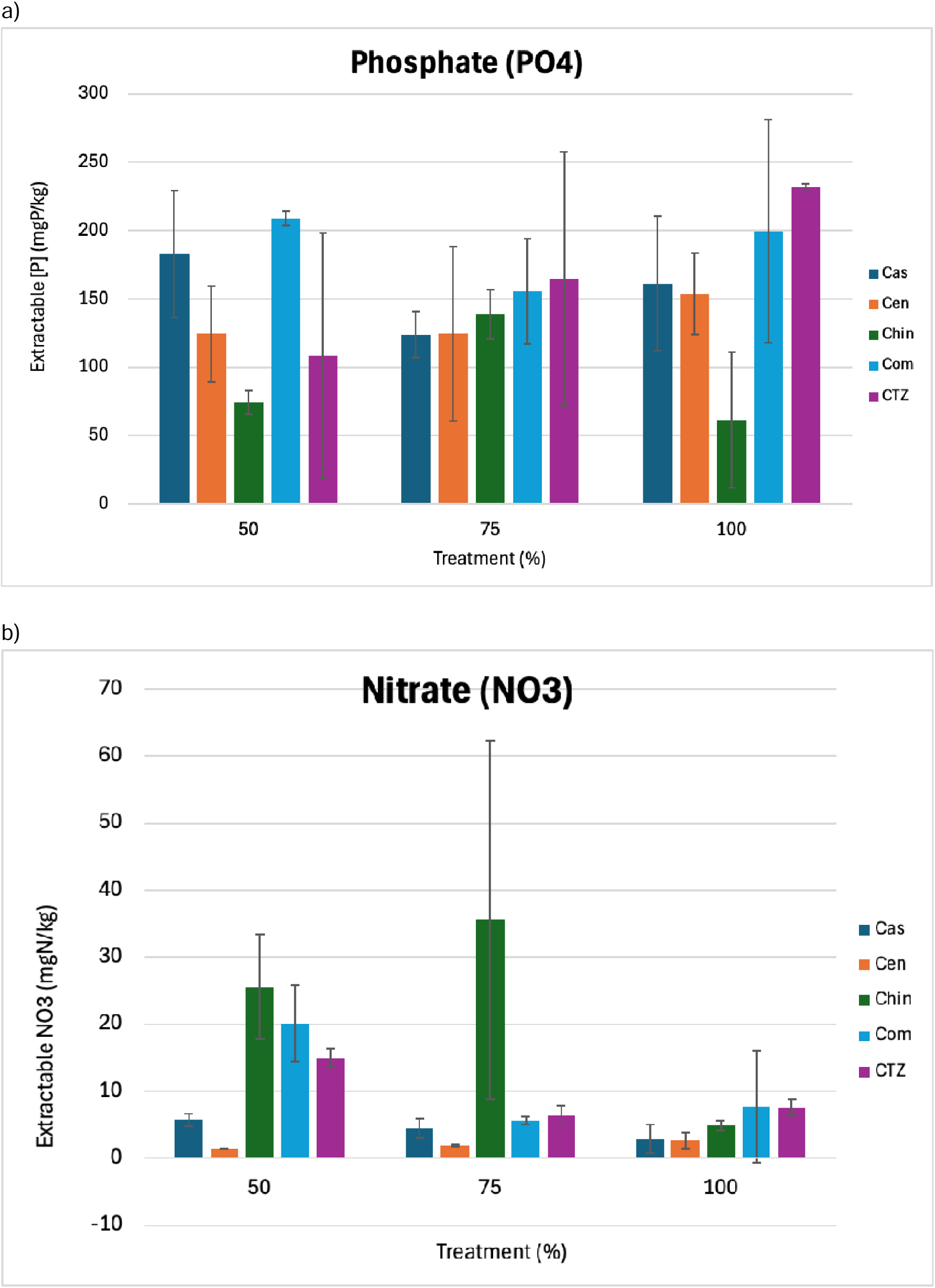
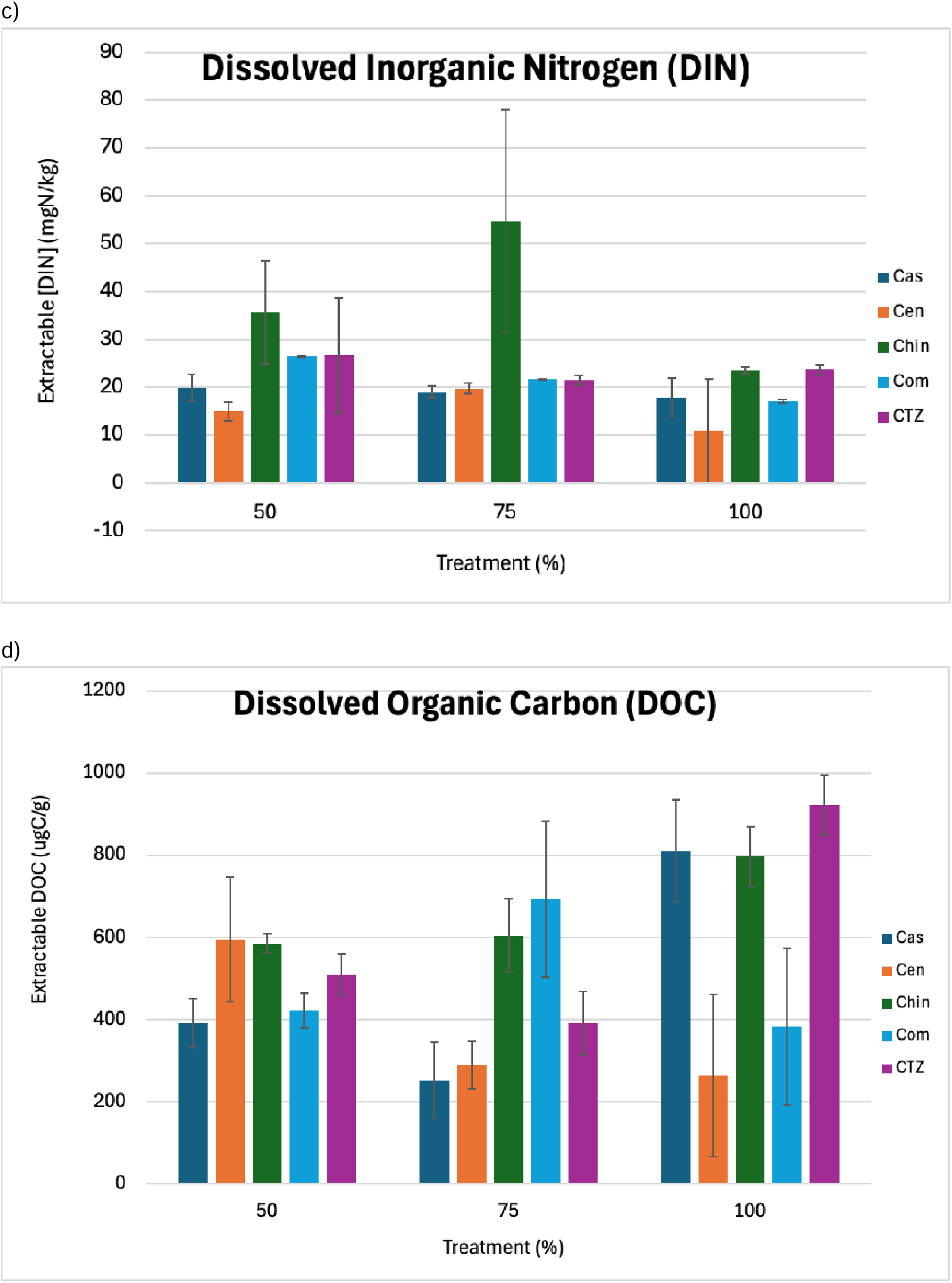
Measured values for extractable nutrient values for phosphate (a), nitrate (b), dissolved inorganic nitrogen (c) and dissolved organic carbon (d) for each cultivar type: Cascade (dark blue), Centennial (orange), Chinook (green), Comet (light blue) and CTZ (purple).

NO3 bioavailability was strongly impacted by both cultivar type (df= 4, F= 14.211, p<0.001) and treatment type (df=2, F= 5.691, p= 0.014) independently and insignificantly when tested for a cultivar∼treatment interaction (df=8, F= 1.548, p= 0.222).

NO3 bioavailability was highest under the moderate drought regime for Chinook and under the severe treatment for Comet and CTZ (Figure 8b). As for Cascade and Centennial, NO3 bioavailability remained relatively consistent between treatment types. Extractable DIN was found to be greatest in the rhizosphere of Chinook cultivars under all three watering regimes, with moderate regime displaying over a 2-fold increase in extractable DIN when compared to the other tested cultivars. Additionally, Comet and CTZ observed the greatest extractable DIN under severe drought conditions (Figure 8c).

Lastly, ANOVA testing of extractable DOC values revealed that cultivar (df= 4, F= 5.576, p= 0.006) and treatment type (df= 2, F= 7.306, p= 0.006) strongly impacted bioavailability of DOC within the rhizosphere. Additionally, an interaction between cultivar type and treatment type was found to strongly impact extractable DOC values (df= 8, F= 7.362, p< 0.001). Cascade, Chinook and CTZ displayed the highest extractable DOC values under the control treatment as well as overall between treatments types (Figure 8d). Under moderate drought conditions, Comet was found to display the greatest extractable DOC values as well as displaying the highest between the three treatment types. Lastly, Centennial was the only cultivar type to display the greatest amount of extractable DOC under severe drought like conditions, nearly doubling values detected for moderate and control conditions.

## Discussion

### American Cultivars Displayed Differential Tolerance to Reduced Watering Strategies

The primary goal of our study was to explore hops’ physiological and rhizospheric responses to reduced watering regimes. As expected, our study revealed that as irrigation levels decreased (moving from control to severe treatment group), phenotypic characteristics such as biomass, weekly growth rate, and leaf shape/size were significantly impacted as a response to the reduced watering regimes. An agricultural product during hops production is the mature hops flower known as the ‘cone’. In order for hops flowers to be produced, plants must complete their vegetative growth phase, achieving a targeted height between 5-6 m in order for ideal yield conditions to occur (Acosta-Ragnel et al., 2021; Rossini et al., 2021). While our study showed that all tested cultivars did become significantly impacted by reduced watering regimes, cultivars such as Chinook, CTZ, and Comet were found to produce greater biomass and greater weekly growth under drought conditions than cultivars like Cascade and Centennial.

Across treatment types, only Comet was found to display no significant differences in biomass production between treatment types, whereas cultivars such as CTZ and Chinook did display some significant differences between treatment types, but overall produced more aboveground biomass than that of Cascade and Centennial. Our ANOVA of weekly growth rates revealed that under the severe drought regime, Comet grew significantly more biomass than Centennial and CTZ, insignificantly more than Cascade (p-value=0.0867) and very similar to that of Chinook. Our ANOVA’s of leaf size and circularity yielded similar patterns as found above. For instance, under ideal watering conditions, Comet and CTZ displayed a significantly larger leaf surface area than that of Cascade but were not significantly larger than Centennial and Chinook. However, under the severe treatment, Comet displayed a significantly larger leaf size than Cascade, Centennial, and Chinook, with CTZ displaying a reduction in leaf size, insignificantly compared to Comet, highlighting the impact of reduced watering strategies. Under the severe drought regime, Comet not only displayed the largest surface area amongst the other four cultivars; additionally, this cultivar saw an increase in surface area compared to that of the control and moderate treatment. Generally, however, leaf sizes began to reduce as water availability decreased, a trend expected for plants under drought stress in order to reduce the plant’s total surface area for water loss via transpiration from photosynthesis (Mohanta et al., 2023).

Similar to leaf size, as drought conditions continue to worsen, it is expected that leaves will become narrower, therefore less circular, as a mechanism to further reduce their surface area to mitigate water loss (Varsamis et al., 2022). Under the control treatment, Comet displayed the lowest leaf circularity score; however, this difference was nonsignificant compared to the cultivars. As water stress increased and water availability decreased, Cascade, Centennial, Chinook, and CTZ generally increased their leaf circularity while Comet remained constant, leading to significantly lower leaf circularity scores between Comet and all other cultivars under the severe drought regime.

### Belowground Associations As Possible Mechanisms for Drought Resiliency

Genetic variation associated with drought response traits may vary phenotypically between cultivars; however, this is not the only possible mechanism for conferring drought tolerance. Belowground associations with rhizospheric microbes may help foster resiliency to drought through mutualistic relationships between crops and their neighboring microbial communities. 16S sequencing of soil samples collected from the rhizospheric region of 30 individuals across our five cultivars and three treatment types revealed that overall, hops rhizospheric communities are dominated by Bacillus, making up 10-30% of the overall bacterial diversity in the region. *Bacillus* composition was found to be on average highest in CTZ and differed significantly between Cascade and Comet. Interestingly, Comet was found to have the lowest percent abundance of this microbe, a trend unexpected due to its unwavering phenotypic resistance in the face of drought.

ANOVA analysis using timepoint (start vs end) as a fixed effect found that 64.5% of the identified bacterial families displayed a significant difference in their percent abundance from the start vs the end of the 12-week experimental period, with nine families displaying highly significant differences (p-value < 0.001). Of these nine families, four families (Bacillaceae, Paenibacillaceae, Planctomycetaceae and Staphylococcaceae) were found to significantly vary in percent abundance between cultivar types. Interestingly, three of these four families are known to contain species of bacteria that have been previously classified as PGPRs, with members of the fourth family, Planctomycetaceae, being shown to improve soil structure, nutrient availability, and nitrogen cycling under periods of osmotic stress, such as drought (Tugbaeva et al., 2025).

Patterns of association between microbial families and cultivar types were also observed. For instance, Chinook and Comet, two cultivars found to display similar phenotypic responses to drought, were also found to be the only two associated with Staphylococcaceae and display similar patterns in abundance with Paenibacillaceae and Planctomycetaceae. Additionally, 16s sequencing of the archaea found within the rhizosphere highlighted that Chinook and Comet displayed the highest abundance of the archaeal family Thaumarchaeota, the only archaea found to be significantly impacted by reduced watering regimes. Abundance of Thaumarchaeota was significantly impacted by watering regimes in Chinook and Comet compared to the remaining cultivars, with abundance highest at the start of the experiment and significantly declining by the end of the 12 weeks.

Interestingly, Thaumarchaeota has been previously reported to confer potential drought tolerance under arid conditions by converting ammonium in the soil to nitrate via nitrification, a substance easily absorbed by the roots and used directly for the proper growth and development of the plant (Burns et al., 2025; Kristy et al., 2022). Data from our experiment revealed that while extractable ammonium was found to not be significantly impacted by either cultivar or treatment type, extractable nitrate varied very significantly between cultivar and treatment types. For Chinook, extractable nitrate was found to be greatest overall under moderate drought regime. Under severe drought conditions however, Chinook still displayed the greatest amount of extractable nitrate compared to other cultivars while Comet displayed an intracultivar best.

Given that Chinook and Comet were two of our cultivars that were found to display the least impact on their growth and development by reduced watering strategies, we hypothesize that microbes presented here, such as Thaumarchaeota, should be further explored for their potential as possible drought conferring mechanisms within the hops rhizosphere. If future studies yield similar findings, potential inoculation treatments may be developed to help aid in conferring drought tolerance, a strategy already being applied to other major cash crops such as maize (Curá et al., 2017; Francioli et al., 2025).

### The Future of Hops Breeding

Production of hops and quality of product are expected to decline over the next 50 years as a response to warming temperatures via climate change (Mozny et al. 2023). In order for breeders to best mitigate these effects, they must first understand the potential response key cultivars may have under these future conditions. Here, we opted to use five of the most cultivated hop cultivars in the United States to assess their potential to withstand reduced watering regimes. Phenotypic data generated from our study suggest that of the five cultivated varieties tested, Comet may serve as the best individual to focus selective breeding tactics towards. We found that under severe drought conditions (50% of the recommended watering), biomass and weekly growth rates did not significantly differ between individuals in the control treatment and the severe treatment, indicating potential resistance to osmotic stress. Additionally, leaf morphological traits such as leaf shape indicated that Comet possesses additional drought-adaptive traits such as narrow leaf shape, a tactic used to minimize leaf surface area and reduce photosynthetic productivity in a time of drought stress.

This begs the question: is Comet the future of hops breeding? The short answer: no. The long answer: It is a potential source of some adaptive variation, but our study revealed an even deeper potential pattern with broader implications. While ANOVAs were performed for each of our tested morphological traits, highlighting the significant differences in Comet’s growth compared to the other four cultivars, a pattern of “grouping” or “response clusters” occurred in the data as well. Throughout our study, we found that Cascade and Centennial displayed similar responses to drought (such as lower biomass, slower growth rates, etc.) to each other than that of Chinook, Comet, and CTZ. Furthermore, in several analyses, Chinook was found to perform very similarly to Comet or, in some cases, better (such as biomass growth under moderate drought conditions) in the face of moderate and severe drought. Additionally, profiling of the rhizospheric communities across our 30 tested cultivars found that Comet and Chinook displayed similar percent abundance for several key PGPR, including being the only two cultivars to foster bacteria from the Staphylococcaceae.

Is this coincidence or something deeper? When we take a step back and look at the pedigree of these two cultivars, we find a commonality: an extensive breeding history of backcrossing with ‘wild American’ breeding lines. In both these cases, their breeding history indicates being backcrossed with wild Utah American hops (USDA Utah 524 and 526). Of the three wild hop varieties known to occur throughout North America, only one species is found in the southwestern United States: *var. neomexicanus*. Coincidentally, this region also typically displays warmer temperatures and lower precipitation patterns than those of its Pacific Northwest counterparts; however, in 50 years from now, climate in this region may be similar to the conditions felt in the future throughout the PNW.

## Conclusion

The primary goal of our study was to evaluate economically important hops and their phenotypic and rhizospheric responses to reduced watering strategies, a very real and near-future possibility. We found that out of the five cultivars we tested, Comet and Chinook consistently performed best under reduced watering strategies, serving as excellent candidates for future breeding efforts. Metabarcoding analysis revealed that hop plants generally harbor several plant growth-promoting rhizobacteria; however, the composition of the rhizosphere significantly differed between tested cultivars, with Comet and Chinook displaying several similarities at the rhizospheric level. Given our findings, we highly suggest that future breeding efforts specifically focus on introgression with wild native American hops populations, specifically of *var. neomexicanus*, in order to remain a competitive player in future hops production.

## Acknowledgments

We would like to thank members of the Sethuraman Lab for their valuable comments on earlier versions of this manuscript. We extend our sincere thanks to Dr. Lluvia Flores-Renteria for permitting use of the SDSU Alvarado Greenhouse. We would also like to thank the Fall 2024 and 2025 Class of BIOL 596-Research Methods in Agricultural Sciences at San Diego State University for their help with data collection for a portion of the growing season.

## Authors Contributions

AMA designed, implemented, collected, analyzed all data, and wrote the manuscript. GV performed all the soil analyses. AS and GV acquired funding and co-wrote the manuscript. CM, KK, ZG helped with all the sample collection, processing, DNA extractions. MG, AA helped with computing the leaf surface area measures. All authors reviewed and approved the final draft.

